# Sequential infection with influenza A virus followed by severe acute respiratory syndrome coronavirus 2 (SARS-CoV-2) leads to more severe disease and encephalitis in a mouse model of COVID-19

**DOI:** 10.1101/2020.10.13.334532

**Authors:** Jordan J. Clark, Rebekah Penrice-Randal, Parul Sharma, Anja Kipar, Xiaofeng Dong, Shaun H. Pennington, Amy E. Marriott, Stefano Colombo, Andrew Davidson, Maia Kavanagh Williamson, David A. Matthews, Lance Turtle, Tessa Prince, Grant L. Hughes, Edward I. Patterson, Ghada Shawli, Daniele F. Mega, Krishanthi Subramaniam, Jo Sharp, Lynn McLaughlin, En-Min Zhou, Joseph D. Turner, Giancarlo Biagini, Andrew Owen, Julian A. Hiscox, James P. Stewart

## Abstract

COVID-19 is a spectrum of clinical symptoms in humans caused by infection with SARS-CoV-2, a recently emerged coronavirus that rapidly caused a pandemic. Coalescence of this virus with seasonal respiratory viruses, particularly influenza virus is a global health concern. To investigate this, transgenic mice expressing the human ACE2 receptor driven by the epithelial cell cytokeratin-18 gene promoter (K18-hACE2) were first infected with IAV followed by SARS-CoV-2. The host response and effect on virus biology was compared to K18-hACE2 mice infected with IAV or SARS-CoV-2 only. Infection of mice with each individual virus resulted in a disease phenotype compared to control mice. Although SARS-CoV-2 RNA synthesis appeared significantly reduced in the sequentially infected mice, they exhibited more rapid weight loss, more severe lung damage and a prolongation of the innate response compared to singly infected or control mice. The sequential infection also exacerbated the extrapulmonary encephalitic manifestations associated with SARS-CoV-2 infection. Conversely, prior infection with a commercially available, multivalent live-attenuated influenza vaccine (Fluenz tetra) elicited the same reduction in SARS-CoV-2 RNA synthesis albeit without the associated increase in disease severity. This suggests that the innate immune response stimulated by infection with IAV is responsible for the observed inhibition of SARS-CoV-2, however, infection with attenuated, apathogenic influenza vaccine does not result in an aberrant immune response and enhanced disease severity. Taken together, the data suggest that the concept of ‘twinfection’ is deleterious and mitigation steps should be instituted as part of a comprehensive public health response to the COVID-19 pandemic.

## Introduction

Coronaviruses were once described as the backwater of virology but the last two decades have seen the emergence of three major coronavirus threats (*1*). First, the emergence of severe acute respiratory syndrome coronavirus (SARS-CoV) in China in 2003. Second, Middle East respiratory syndrome coronavirus (*2*) in Saudi Arabia in 2012 and now SARS-CoV-2 originating in China in 2019. Whilst SARS-CoV was eradicated, both MERS-CoV and SARS-CoV-2 represent current ongoing health threats, and a greater understanding is required to develop robust interventions for future emergent coronaviruses. Coronaviruses share similar genome architectures and disease profiles and generally cause respiratory and gastrointestinal illnesses (*1*). However, some animal/avian coronaviruses can also affect other organ systems, causing, for example, demyelination and nephritis. The sheer scale of the COVID-19 outbreak has highlighted hitherto unexpected aspects of coronavirus infection in humans, including long term disease complications once the virus has been cleared.

Infection of humans with SARS-CoV-2 results in a range of clinical courses, from asymptomatic to severe infection and subsequent death in not only at-risk individuals but also a small proportion of otherwise healthy individuals across all age groups. Severe infection in humans is typified by cytokine storms (*3, 4*), pneumonia and kidney failure. Examination of post-mortem tissue reveals a disconnect between viral replication and immune pathology (*5*). A range of other clinical signs also occur, including gastrointestinal symptoms such as vomiting, diarrhoea, abdominal pain and loss of appetite as well as loss of taste and smell (anosmia). A small number of patients have no overt respiratory symptoms at all. Typically, patients with severe COVID-19 present to hospital in the second week of illness. There is often a precipitous decline in respiratory function, without necessarily much in the way of “air hunger.” Once intubated, these patients have unique ventilatory characteristics, where they can be ventilated with relatively low inspired oxygen concentrations but need high positive end expiratory pressures.

Respiratory infections in humans and animals can also be synergistic in which an initial infection can exacerbate a secondary infection or vice versa. When multiple pathogens are in circulation at the same time this can lead to cooperative or competitive forms of pathogen-pathogen interactions (*6*). This was evident during the 1918 Spanish influenza A virus outbreak (IAV) where secondary bacterial pneumonia was thought to be a leading cause of death (*7*). Coinfections in other viral diseases, such as in patients with Ebola virus disease, have also been shown to contribute to the host response and outcome (*8*). Global influenza cases decreased due to the lockdowns implemented to contain SARS-CoV-2 spread (*9–11*), the lifting of these lockdowns in 2021 and 2022 has resulted in the return of seasonal influenza outbreaks (Weekly U.S. Influenza Surveillance Report, CDC, https://www.cdc.gov/flu/weekly/index.htm). With ongoing the SARS-CoV-2 waves caused by emerging variants of concern (VOCs) and a return to seasonal IAV outbreaks, coinfection with these respiratory pathogens is likely and this may exacerbate clinical disease and potentially outcome.

Previous work has shown coinfections are present in patients with severe coronavirus infection. For SARS-CoV co-circulation of human metapneumovirus was reported in an outbreak in Hong Kong. However, data suggested that outcomes were not different between patients with identified coinfections and those with SARS-CoV alone (*12*). For MERS-CoV, four cases of coinfection with IAV were described, and although no data was presented on the severity of symptoms this sample size would be too small to allow any meaningful conclusions (*13*). Post-mortem studies from patients with COVID-19 in Beijing (n=85) identified IAV in 10% of patients, influenza B virus in 5% and respiratory syncytial virus (RSV) in 3% of patients, but the absence of a carefully selected control arm prohibits conclusions to be drawn (*14*). Additionally, there have been several case reports of coinfections with IAV and SARS-CoV-2 in humans with severe outcomes (*15–20*) with one study from the UK reporting that patients with a coinfection exhibited a ∼6 times higher risk of death(*21*). Whilst this suggests that coinfection is synergistic, this study also found that the risk of testing positive for SARS-CoV-2 was 68% lower among individuals who were positive for IAV infection, implying that the two viruses may competitively exclude each other(*21*).

Whilst the analysis of post-mortem tissue is extremely informative in what may have led to severe coronavirus infection and death, the analysis of the disease in severe (but living cases) is naturally restricted by what tissues can be sampled (e.g. blood, nasopharyngeal swabs and bronchioalveolar lavages). Therefore, animal models of COVID-19 present critical tools to fill knowledge gaps for the disease in humans and for screening therapeutic or prophylactic interventions. Compatibility with a more extensive longitudinal tissue sampling strategy and a controlled nature of infection are key advantages of animal models (*22*). Studies in an experimental mouse model of SARS-CoV showed that coinfection of a respiratory bacterium exacerbated pneumonia (*23*). Different animal species can be infected with wild-type SARS-CoV-2 to serve as models of COVID-19 and these include mice, hamsters, ferrets, rhesus macaques and cynomolgus macaques. The K18-hACE2 transgenic (K18-hACE2) mouse, where hACE2 expression is driven by the epithelial cell cytokeratin-18 (K18) promoter, was developed to study SARS-CoV pathogenesis(*24*). This mouse is being used as a model that mirrors many features of severe COVID-19 in humans to develop understanding of the mechanistic basis of lung disease and to test pharmacological interventions(*25, 26*).

IAV and SARS-CoV-2 co- and sequential infections have already been studied in hamsters, with partly controversial results. In a first study, simultaneous or subsequent infection 24 hours apart, they led to more severe pneumonia (*27*). A similar study then found evidence that co-infection with SARS-CoV-2 leads to more widespread IAV infection in the lungs (*28*), while a third study described impaired IAV replication in lungs when hamsters were infected early and 10 days after SARS-CoV-2 inoculation, but no effect when IAV infection took place on day 21 post SARS-CoV-2 infection (*29*). With the liklihood of flu seasons concomitant with waves of SARS-CoV-2 infections there is an obvious public health concern about the possibility of enhanced morbidity and mortality in coinfected individuals. The aim of this work was to use an established pre-clinical model of COVID-19 to study the consequences of coinfection with SARS-CoV-2 and IAV, defining the associated clinical, pathological and transcriptomic signatures.

## Results

### Sequential infection with pathogenic IAV, but not Fluenz Tetra plus SARS-CoV-2 leads to enhanced disease

To assess how coinfection with influenza virus affected SARS-CoV-2 infection, the established K18-hACE2 mouse model of SARS-CoV-2 was utilised (*24*). We used a clinical isolate of SARS-CoV-2 (strain hCoV-19/England/Liverpool_REMRQ0001/2020)(*30*). Importantly, sequence of the virus stock demonstrated that this isolate did not contain deletions or mutations of the furin cleavage site in the S protein (*31*). A schematic of the experimental design is shown in Fig. 1A. Four groups of mice (n = 8 per group) were used. At day 0, two groups were inoculated intranasally with 10^2^ PFU IAV (strain A/X31) and two groups with PBS. After three days, two groups were inoculated intranasally with 10^4^ PFU of SARS-CoV-2. This generated four experimental groups: Control, IAV only, SARS-CoV-2 and IAV + SARS-CoV-2 only (Fig. 1B). Control mice maintained their body weight throughout. Mice infected with IAV displayed a typical pattern of weight loss, reaching a nadir (mean 17% loss) at 7 dpi before starting recovery. SARS-CoV-2-infected animals started to lose weight at day 7 (4 dpi) and carried on losing weight up to day 10 (mean 15% loss). Mice infected with IAV then SARS-CoV-2 had a significantly accelerated weight loss as compared with IAV-infected mice from day 4; this was most severe at day 6 (mean 19%), followed by a recovery to day 8 (mean 14% loss) before losing weight again (mean 17% loss) (Fig. 2A). As well as accelerated weight loss, IAV + SARS-CoV-2-infected mice exhibited more severe respiratory signs and a significantly more rapid mortality (assessed by a humane endpoint of 20% weight loss) as compared with mice infected with either virus alone (Fig. 2B). To investigate the effect of infecting mice with a non-pathogenic vaccine formulation of influenza, K18-ACE2 mice were dosed intranasally with Fluenz Tetra 3 days prior to SARS-CoV-2 infection (Fig. 1B). Mice that received Fluenz Tetra only, showed no significant deviation in weight loss in comparison to the PBS control (Fig 2A). Mice that received Fluenz Tetra immunisation followed by a SARS-CoV-2 infection initially had a slight reduction in weight at day 1 and day 2, however, between day 3 and day 8 resembled the PBS control mice, although at day 10 two coinfected mice had lost 13% and 9% of their body weight while two continued to gain weight (Fig. 2A). The animals showing weight loss did not exhibit severe respiratory signs associated with SARS-CoV-2 infection or coinfection.

**Figure 1.**
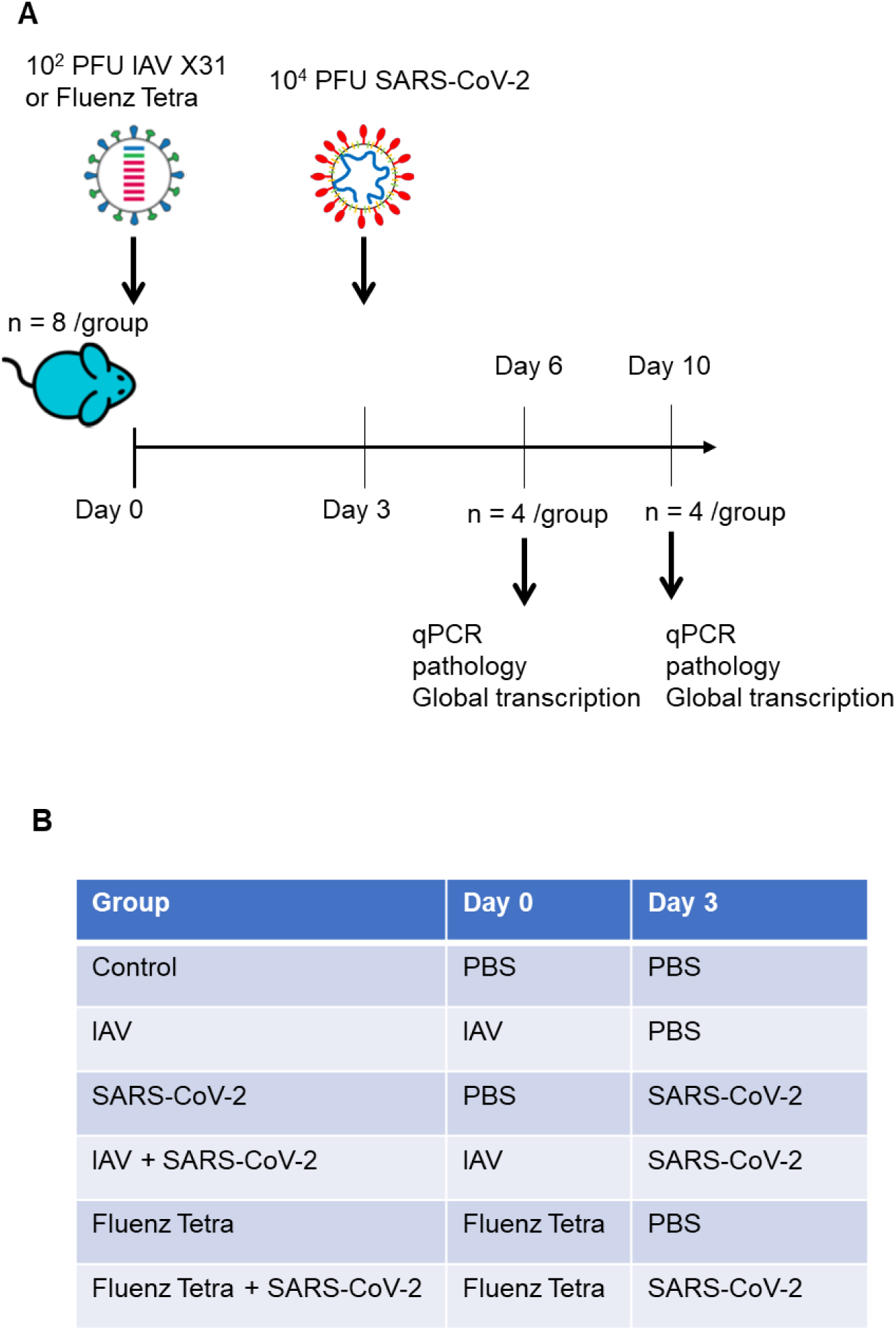
**A)** Schematic diagram of the experimental design for infection of K18-hACE2 mice sequentially with IAV strain A/X31 and SARS-CoV-2 (hCoV-19/England/Liverpool_REMRQ0001/2020). **B)** Table showing exposures given to the four individual groups of mice.

**Figure 2.**
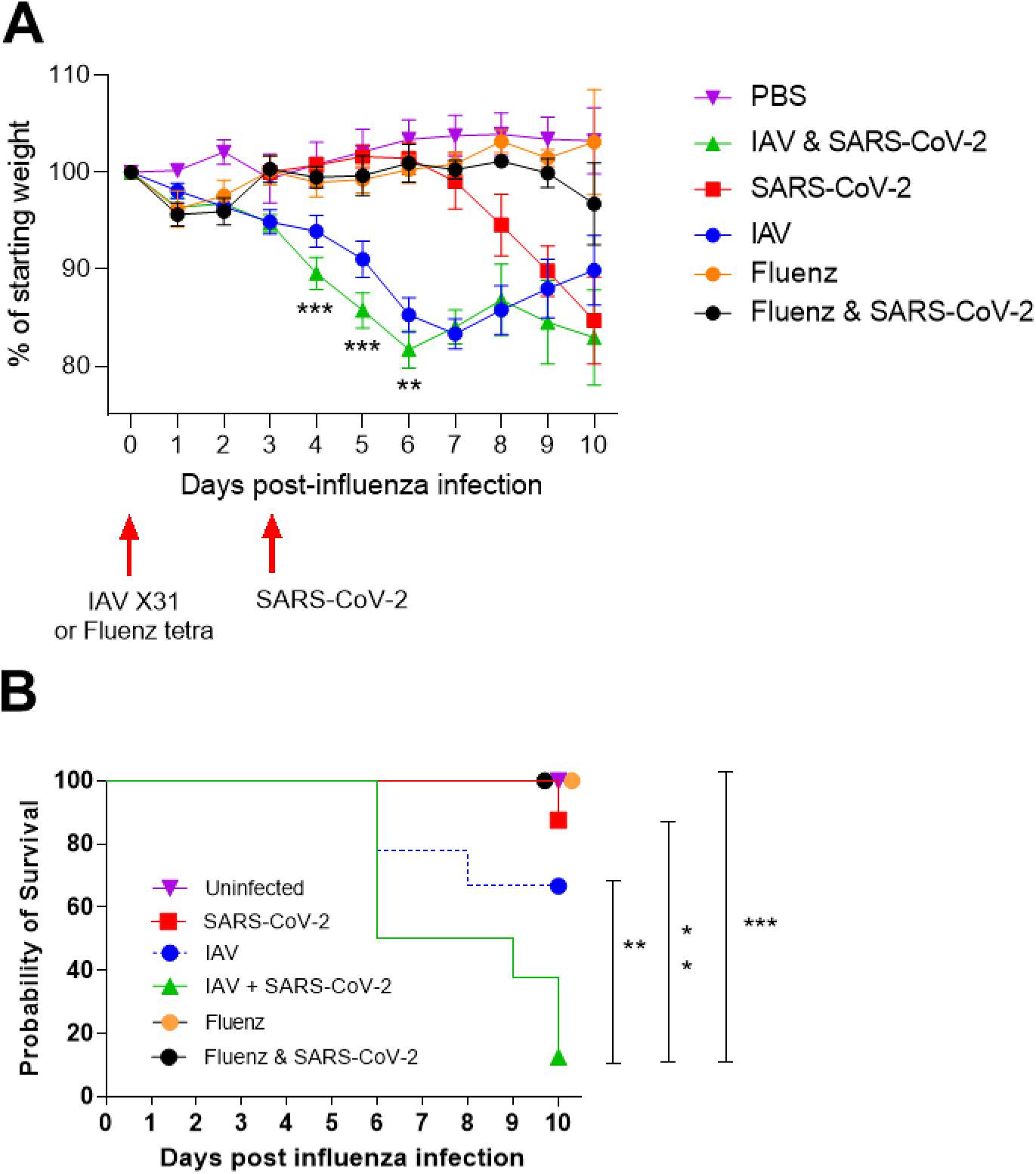
Coinfection with IAV and SARS-CoV-2 leads to enhanced weight loss and more rapid mortality. K18-hACE2 mice were challenged intranasally with IAV strain X31 (10^2^ PFU) or immunized with Fluenz Tetra, and challenged 3 days later with 10^4^ PFU SARS-CoV-2. **A)** Mice were monitored for weight loss at indicated time points (n = 8). Data represent the mean value ± SEM. Comparisons were made using a repeated-measures two-way ANOVA (Bonferroni post-test). **B)** Survival was assessed at indicated time points (n = 8). Comparisons were made using log-rank (Mantel-Cox) test. ** represents P < 0.01; *** represents P < 0.001

### Coinfection of SARS-COV-2 and IAV results in reduced SARS-COV-2 viral load at day 6 but not day 10 post IAV

In order to determine whether the coinfection of SARS-CoV-2 and IAV was cooperative or competitive total RNA was extracted from the lungs of the K18-hACE2 mice and viral loads were quantified using qRT-PCR. At day 6 (3 dpi), the SARS-CoV-2 infected mice exhibited 10,000-fold higher levels of viral load than at day 10 (7 dpi) (mean 6 × 10^12^ vs 2.8 × 10^8^ copies of N/µg of RNA) indicating that peak viral replication takes place before the onset of clinical signs at 4 dpi (Fig. 3A). At this time point the mice infected with SARS-CoV-2 alone displayed significantly higher levels of viral RNA than the mice with IAV and SARS-CoV-2 coinfection (mean 6 × 10^12^ vs ∼2 × 10^9^ copies of N/µg of RNA) (Fig. 3A). The mice preimmunised with Fluenz Tetra exhibited reduced levels of viral RNA compared to both SARS-CoV-2 singly and IAV coinfected mice, with 2/4 mice exhibiting no detectable viral RNA. However, by day 10 the SARS-CoV-2 and IAV X31 coinfected and singly infected mice exhibited nearly identical levels of SARS-CoV-2 RNA (mean 2 × 10^8^ vs 8.1 × 10^8^ copies of N/µg of RNA) (Fig. 3A). Conversely, SARS-CoV-2 RNA levels were undetectable in 2/4 of the mice pre-immunized with Fluenz Tetra, with only one animal displaying similar levels of viral RNA to the SARS-CoV-2 singly and IAV coinfected animals. The levels of infectious virus generally corresponded with the copies of N RNA, except at day 10, when there was no infectious virus in mice infected with SARS-CoV-2 alone whereas the level of infectious virus in coinfected mice was similar at both day 6 and day 10 (10^2^ PFU/lung) (Fig. 3C). Conversely, at day 6, the mice infected with IAV alone showed similar levels of IAV RNA compared to the coinfected mice (mean 1.3 × 10^7^ vs 1 × 10^7^ copies of M/µg of RNA) and by day 10 both the singly infected mice and coinfected mice did not display any detectable IAV RNA, demonstrating similar levels of IAV clearance (Fig. 3D). Fluenz Tetra immunized mice displayed reduced levels of IAV viral RNA compared to IAV infected mice (mean 5 × 10^3^), in line with the reduced replication expected of attenuated IAV. To investigate viral replication qPCR was employed to quantify viral subgenomic mRNA (sgRNA) transcripts. Unlike viral genomes, sgRNAs are not incorporated into virions, and can therefore be utilised to measure active virus infection. The amount of sgRNA in the SARS-CoV-2 infected mice was concomitant with the viral load, appearing to be 100-fold higher at day 6 (3dpi) than day 10 (7dpi) (mean 6.2 × 10^6^ vs 5.4 × 10^4^ copies of E sgRNA/µg of RNA) (Fig. 3B). Similarly, the amount of sgRNA was significantly lower in the coinfected mice compared to the SARS-CoV-2 singly infected mice (mean 6.2 × 10^6^ vs 1.7 × 10^4^ copies of E sgRNA/µg of RNA) however, by day 10 (7dpi) both coinfected and singly infected mice displayed similar levels of sgRNA (mean 5.4 × 10^4^ vs 3.5 × 10^5^ copies of E sgRNA/µg of RNA) (Fig. 3B). Fluenz tetra immunized mice exhibited further reduced levels of sgRNA at day 6, with only one mouse exhibiting detectable sgRNA, however by day 10 2/4 mice displayed detectable sgRNA.

**Figure 3.**
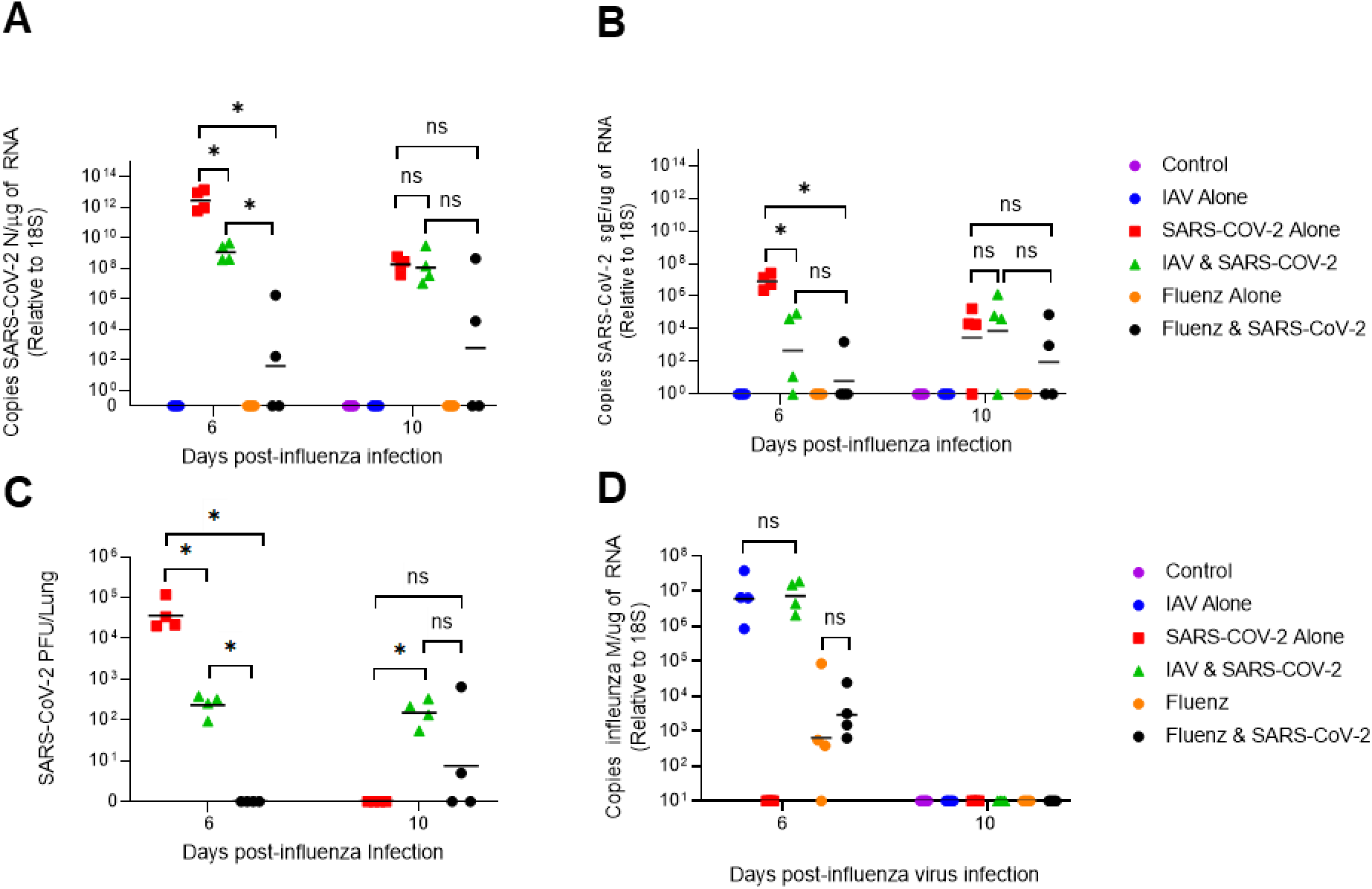
Viral loads and SARS-CoV-2 sgRNA levels in single and coinfected mice. K18-hACE2 mice were challenged intranasally with IAV strain X31 (10^2^ PFU) and 3 days later with 10^4^ PFU SARS-CoV-2 (n = 4). RNA extracted from lungs was analysed for virus levels by qRT-PCR. Assays were normalised relative to levels of 18S RNA. Data for individual animals are shown with the median value represented by a black line. **A)** SARS-CoV-2 viral load was determined using qRT-PCR for the N gene**. B)** Levels of SARS-CoV-2 sub-genomic RNA (sgRNA) for the E gene. **C)** SARS-CoV-2 titre was determined by plaque assay on Vero E6 cells. **D)** IAV load was determined using RT-PCR for the M gene. Side-by-side comparisons were made using Mann-Whitney U test. * represents p < 0.05

### Coinfection leads to complementary and enhanced pathological processes

Transgenic mice carrying the human ACE2 receptor under the control of the keratin 18 promoter (K18-hACE2) have been widely used as a COVID-19 model (*25*). As a basis for the assessment of the effect of IAV and SARS-CoV-2 in these mice, a histological examination of major organs/tissues was performed. This confirmed that the transgenic approach had not resulted in phenotypic changes. Comparative staining of wild type and K18-hACE2 mice for ACE2, using an antibody against human ACE2 that also cross-reacts with mouse ACE2, also confirmed that transgenesis had not altered the ACE2 expression pattern: in the lung, ACE2 was found to be expressed by respiratory epithelial cells and very rare type II pneumocytes (Supplementary Fig. S1A, B). Expression was also seen in the brain microvasculature where it has recently been shown to be expressed specifically by the pericytes (*32*)(Supplementary Fig. S2 A2, A3) and liver sinusoids and in renal tubular epithelial cells. The expression was not substantially affected by either viral infection (Supplementary Fig. S1C-F; Fig. S2 B, C).

At 6 days post IAV infection, the transgenic mice exhibited the pulmonary changes typically seen in wild type mice after IAV X31 infection at this time point. We observed epithelial cell degeneration and necrosis in several bronchioles which also contained debris in the lumen (Fig. 4B). There were occasional small focal peribronchial areas where alveoli also exhibited necrotic cells (Fig. 4B). IAV antigen was found in epithelial cells in bronchi and bronchioles, in type I and II pneumocytes in affected alveoli, and in few randomly distributed individual type II pneumocytes (data not shown). Vessels showed evidence of lymphocyte recruitment, vasculitis, and perivascular lymphocyte infiltration. Comparative assessment of the lungs in wild type mice at the same time point post infection confirmed that the genetic manipulation indeed had no effect on the response of mice to IAV infection (data not shown). At the comparative time point, SARS-CoV-2 single infection (day 6, 3 dpi) was associated with mild changes, represented by a mild increase in interstitial cellularity, evidence of type II pneumocyte activation (Fig. 4C, 5A), occasional desquamated alveolar macrophages/type II pneumocytes and single erythrocytes in alveolar lumina, and a multifocal, predominantly perivascular mononuclear infiltration with recruitment of leukocytes into vascular walls (vasculitis) (Fig. 4D). Infiltrating cells were predominantly macrophages, with T cells mainly around vessels and a low number of disseminated B cells (Fig. 5); macrophages and T cells were also found to emigrate from veins (Fig. 5D, E). Viral antigen was found in multifocal patches of individual to large groups of alveoli, randomly distributed throughout the parenchyma, within type I and type II pneumocytes (Fig. 5C), but not within bronchiolar epithelial cells (Fig. 5B). Double infection at this time point, i.e. 6 days after IAV infection and 3 days after SARS-CoV-2 infection, was associated with histological changes almost identical to those induced by IAV, although they appeared to be slightly more extensive (Fig. 4E, F). IAV antigen expression had a distribution and extent similar to that seen in single IAV infection at the same time point. It was observed in epithelial cells in bronchi and bronchioles, in type I and II pneumocytes in affected alveoli, and in few randomly distributed individual type II pneumocytes (Fig. 6B). SARS-CoV-2 expression was less intense than in SARS-CoV-2-only infected mice. Viral antigen was observed in random individual or small groups of alveoli (Fig. 6C), in type I and II pneumocytes (Fig. 6C inset).

**Figure 4.**
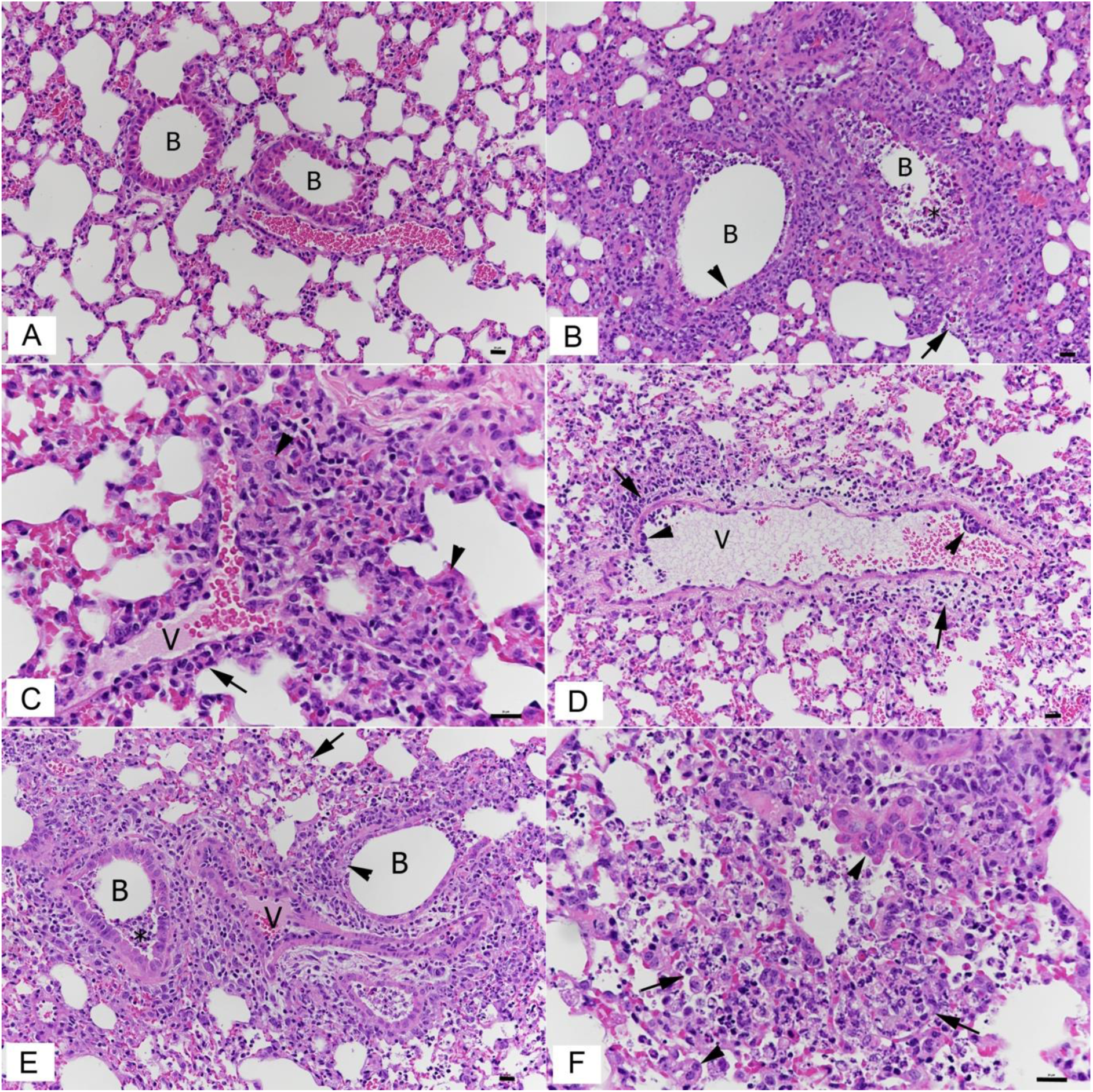
Lungs, K18-hACE2 transgenic mice, after mock infection or at day 6 post infection with IAV and day 3 post infection with SARS-CoV-2 in single and double infections. **A)** Mock infected control animal. Normal lung. **B)** IAV infected animal; 6 dpi. Bronchioles (B) exhibit necrosis (arrowhead) of a variable number of epithelial cells and contain degenerate cells in the lumen (*). The parenchyma adjacent to affected bronchioles often exhibits individual alveoli with necrotic epithelial cells (arrow). **C, D)** SARS-CoV-2 infected animal; 3 dpi. The parenchyma exhibits multifocal activation of type II pneumocytes (C: arrowheads), and there is evidence of vasculitis, represented by leukocyte infiltration of vessel (V) walls (D: arrowheads) and perivascular infiltrates (arrows). **E, F)** IAV (6 dpi) and SARS-CoV-2 (3 dpi) double infection. The IAV associated changes, with necrosis of bronchiolar epithelial cells (E: arrowhead), debris in bronchiolar lumina (*), focal necrosis of alveolar epithelial cells (arrows) as well some activation and hyperplasia of type II pneumocytes (F: arrowheads), dominate the histological picture. B – bronchiole; V – vessel. HE stain; Bars represent 20 µm.

**Figure 5.**
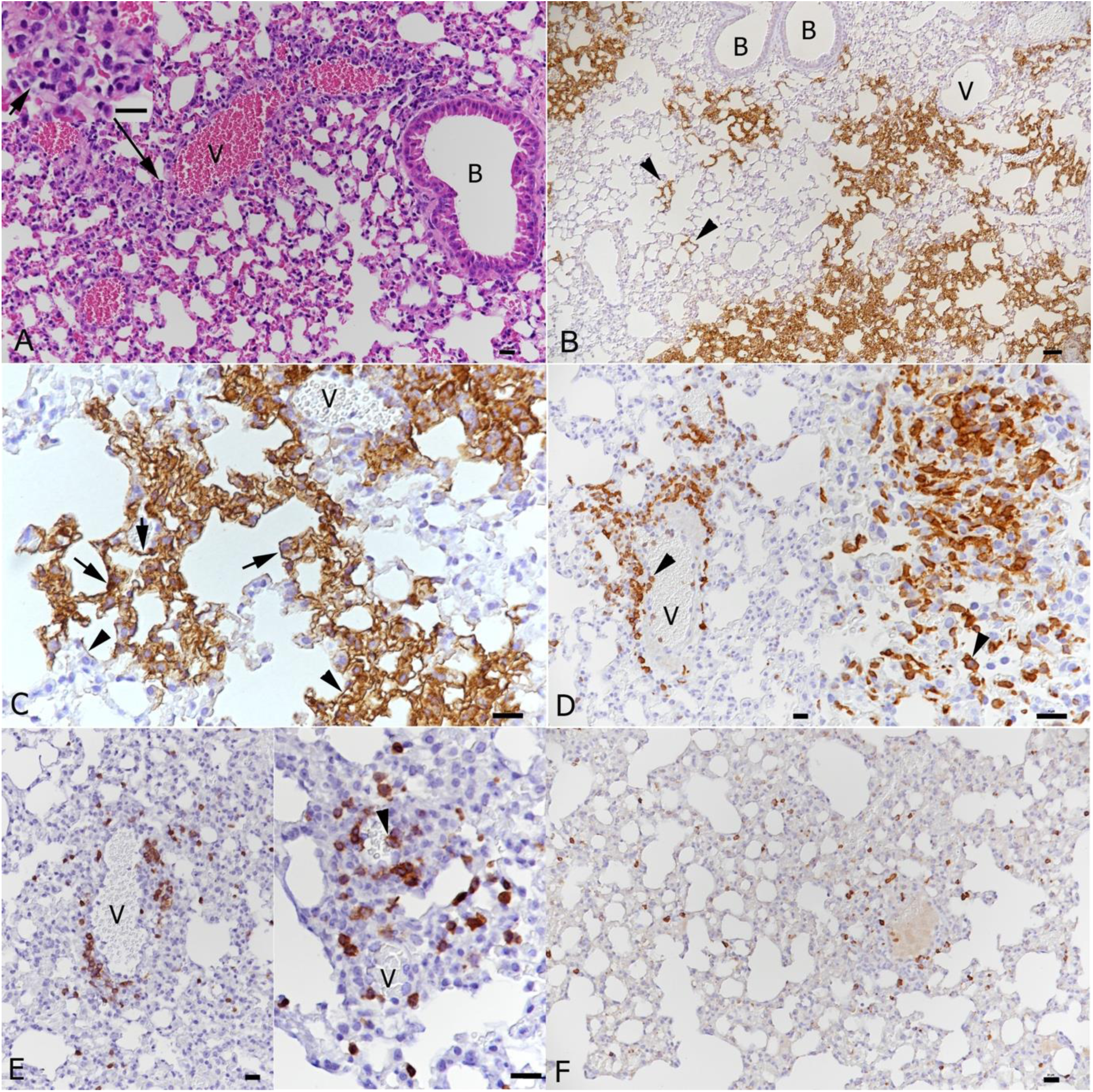
Lungs, K18-hACE2 transgenic mice, at day 3 post infection with SARS-CoV-2. **A)** There is mild perivascular mononuclear infiltration, and the parenchyma exhibits mild multifocal activation of type II pneumocytes (inset). **B, C)** Staining for SARS-CoV-2 antigen reveals random multifocal areas of SARS-CoV-2 infection, affecting both individual alveoli (B: arrowheads) and large parenchymal areas. Viral antigen expression is seen in type I pneumocytes (C: arrowheads) and type II pneumocytes (C: arrows). **D)** Staining for macrophages (Iba1+) shows recruitment from (left image, arrowhead: monocytes attached to the endothelium of a vein) and accumulation of monocytes around veins, macrophage accumulation in the parenchyma and desquamation of alveolar macrophages (right image, arrowhead). **E)** T cells (CD3+) are less numerous than macrophages and are mainly found in the perivascular infiltrates. They are also recruited from the blood (right image, arrowhead). **F)** B cells (CD45R/B220+) are seen in low numbers, and disseminated in the parenchyma. B - bronchiole; V - vessel. HE stain; immunohistology, hematoxylin counterstain. Bars represent 20 µm.

**Figure 6.**
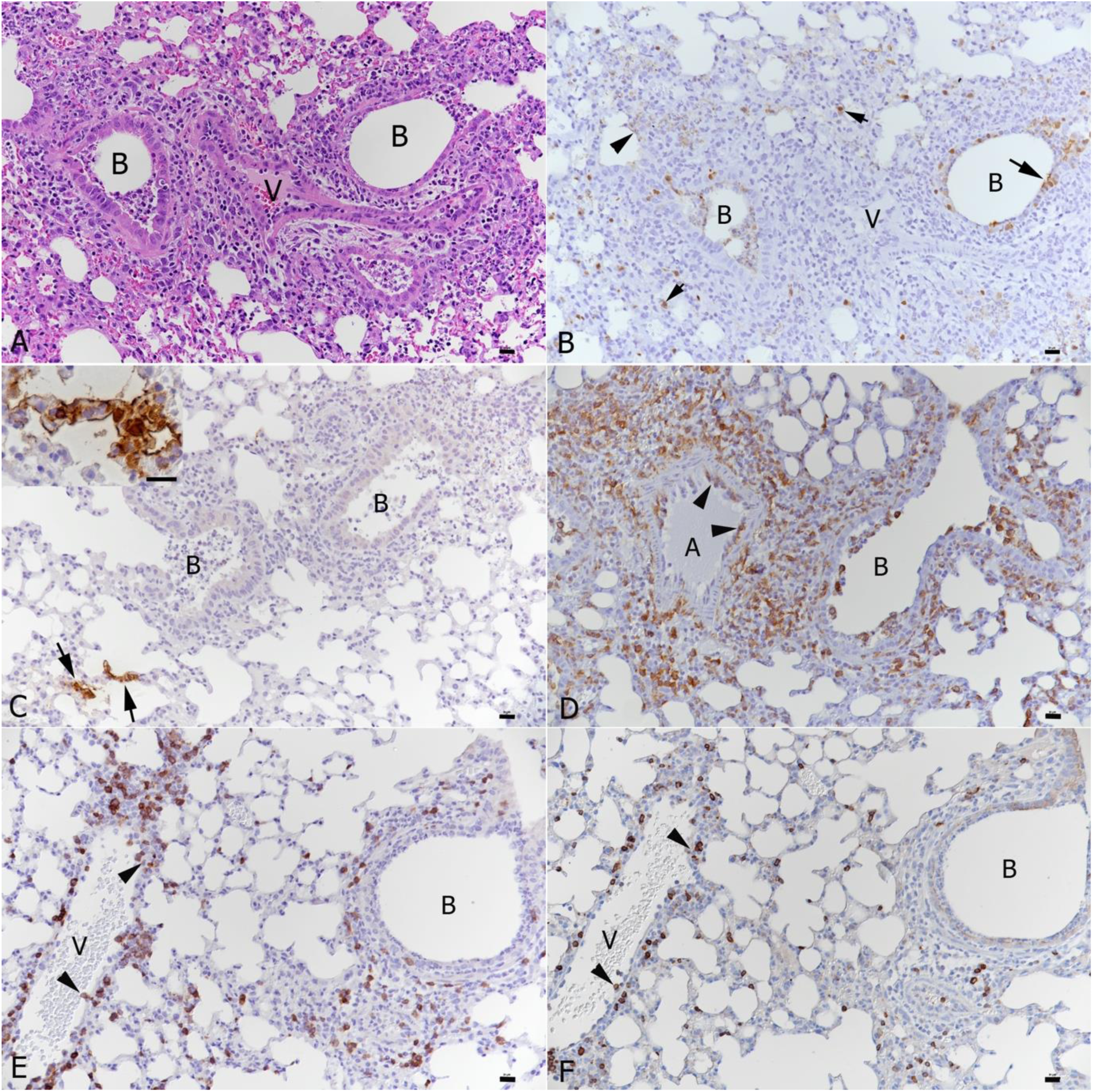
Lungs, K18-hACE2 transgenic mice, at day 6 post infection with IAV and day 3 post infection with SARS-CoV-2 in double infection. **A)** The IAV associated changes dominate (see also Fig. 4E). **B)** This is confirmed by staining for IAV antigen which is detected in bronchiolar epithelial cells (arrow), occasional type I pneumocytes (arrowhead) and disseminated type II pneumocytes (short, small arrows). **C)** SARS-CoV-2 infection is seen in areas not affected by IAV induced changes (B: bronchioles with IAV changes) and mainly in individual alveoli where both type I and type II pneumocytes are found to express viral antigen (inset). **D)** Macrophages (Iba1+) are abundant around affected bronchioles and in the exudate in the bronchiolar lumen and are recruited from the blood into the perivascular infiltrates (arrowheads: rolling and emigrating monocytes). **E)** T cells (CD3+) are recruited in moderate numbers from the blood (arrowheads) into the perivascular infiltrates. **F)** B cells (CD45R/B220+) are recruited in low numbers from the blood (arrowheads) into the perivascular infiltrates. B - bronchiole; V - vessel. HE stain; immunohistology, hematoxylin counterstain. Bars represent 20 µm.

Four days later, at the endpoint of the experiment, i.e. at 10 days after IAV infection and 7 days of SARS-CoV-2 infection, the histopathological features had changed. Single IAV infection had by then almost entirely resolved, however, the lungs exhibited changes consistent with a regenerative process, i.e. mild to moderate hyperplasia of the bronchiolar epithelium with adjacent multifocal respiratory epithelial metaplasia/type II pneumocyte hyperplasia, together with mild to moderate lymphocyte dominated perivascular infiltration (Fig. 6A). Interestingly, the hyperplastic epithelium was found to lack ACE2 expression (Supplementary Fig. S1E). At this stage, the effect of SARS-CoV-2 infection was more evident. Single infection had resulted in multifocal areas with distinct type II pneumocyte activation and syncytial cell formation (Fig. 7B), mononuclear infiltration and mild to moderate lymphocyte-dominated vasculitis and perivascular infiltration. There were also a few focal areas of mild desquamative pneumonia with intra-alveolar macrophages/type II pneumocytes, edema and fibrin deposition (Fig. 7C). Macrophages and T cells dominated in the infiltrates (Fig. 7D, E), whereas B cells were found disseminated in low numbers (Fig. 7F). The SARS-CoV-2 associated changes were also observed in the double infected mice (Fig. 8C-F) where they were generally more pronounced (Fig. 8B, C) and present alongside equally pronounced regenerative changes attributable to IAV infection (moderate hyperplasia of the bronchiolar epithelium with adjacent multifocal respiratory epithelial metaplasia/type II pneumocyte hyperplasia; Fig. 8A). Also, in this group of animals, macrophages were the dominant infiltrating cells. However, the number of T cells was comparatively low (Fig. 8D, E). B cells were generally rare, but occasionally formed small aggregates in proximity to areas of type II pneumocyte hyperplasia (Fig. 8F). Interestingly, the pattern of viral antigen expression had not changed with time; it was detected in type I and II pneumocytes of unaltered appearing alveoli (data not shown).

**Figure 7.**
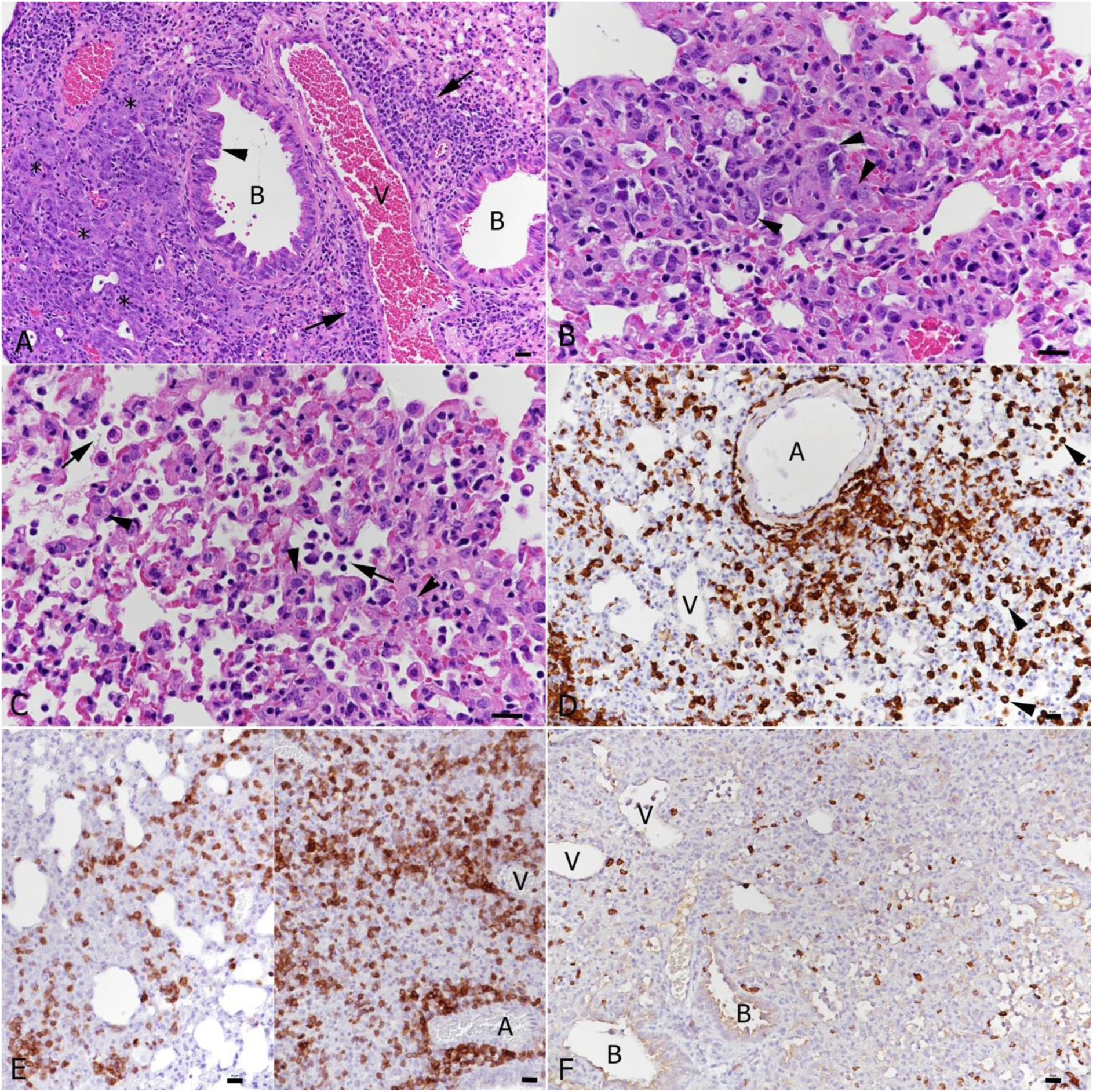
Lungs, K18-hACE2 transgenic mice, at day 10 post infection with IAV and day 7 post infection with SARS-CoV-2 in single infections. **A)** IAV infected animal; 10 dpi. Bronchioles (B) exhibit epithelial cell hyperplasia (arrowhead) and there is type II pneumocyte hyperplasia (*) in the adjacent parenchyma. Vessels (V) exhibit variably intense lymphocyte-dominated perivascular infiltrates (arrows). **B-F**. SARS-CoV-2 infected animal; 7 dpi. **B)** There are abundant activated type II pneumocytes which also show syncytia formation (arrowheads). **C)** There are also focal changes consistent with acute pneumonia, with desquamation of alveolar macrophages/type II pneumocytes (arrows) and type II pneumocyte activation (arrowheads). **D)** Macrophages (Iba1+) are abundant in perivascular infiltrates and within the altered parenchyma. **E)** T cells (CD3+) are also abundant in the parenchymal infiltrates. **F)** B cells (CD45R/B220+) are seen in low numbers disseminated in the parenchymal infiltrates. A - artery; B - bronchiole; V - vessel. HE stain; immunohistology, hematoxylin counterstain. Bars represent 20 µm.

**Figure 8.**
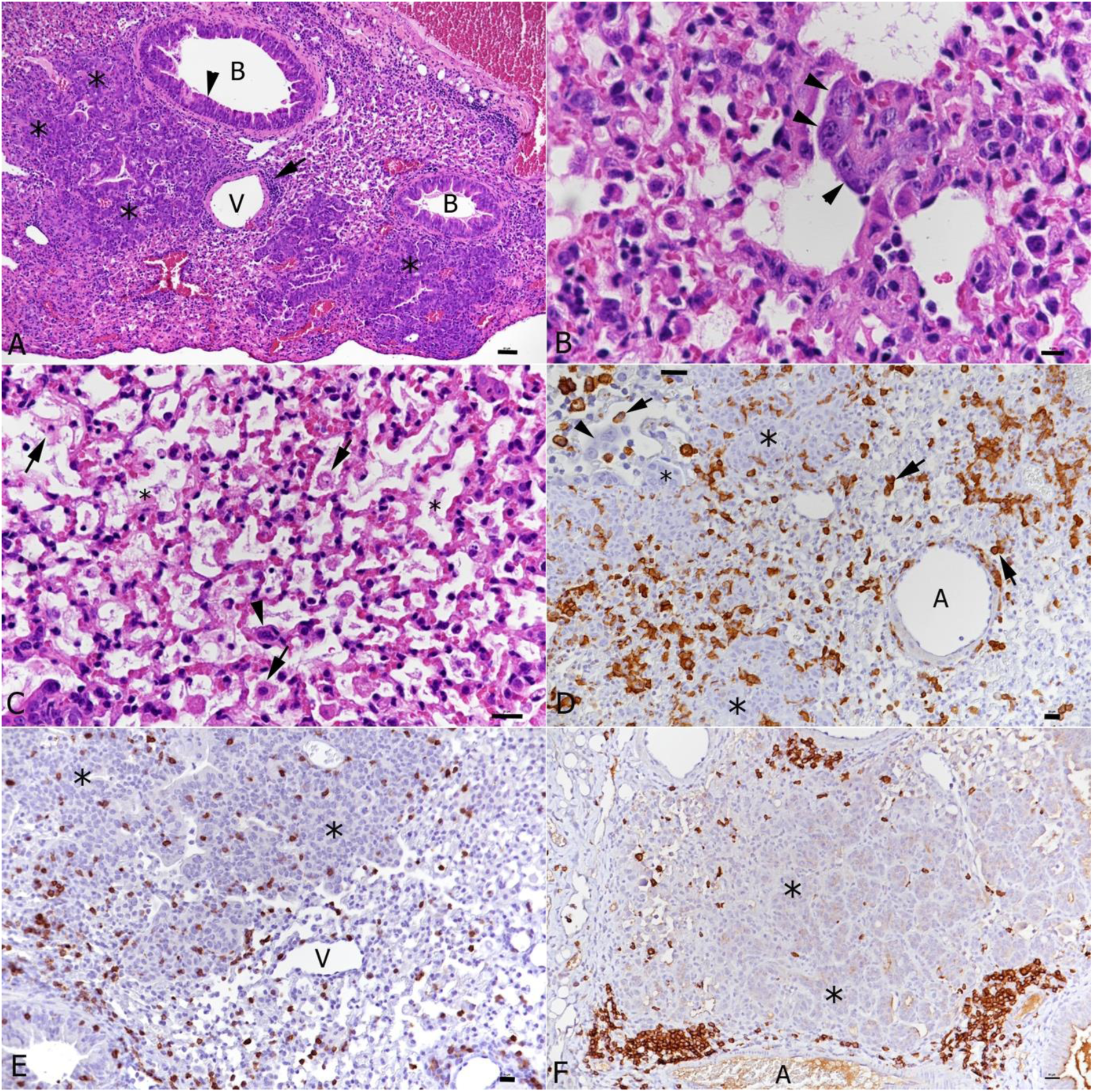
Lungs, K18-hACE2 transgenic mice, at day 10 post infection with IAV and day 7 post infection with SARS-CoV-2 in double infection. **A)** There are abundant changes consistent with those seen in single IAV infected mice, i.e. epithelial cell hyperplasia in bronchioles (B), multifocal type II pneumocyte hyperplasia (*), and perivascular lymphocyte dominated infiltrates (arrow). **B, C)** Changes attributable to SARS-CoV-2 infection. These comprise type II pneumocyte activation and syncytia formation (B: arrowheads) and acute pneumonia (C), with desquamation of alveolar macrophages/type II pneumocytes (arrows) and type II pneumocyte activation (arrowheads). In more severe cases, alveoli occasionally contain fibrin and hyaline membranes (*). **D)** Macrophages (Iba1+) form focal parenchymal infiltrates and are found around areas of type II pneumocyte hyperplasia (*). There are also desquamated alveolar macrophages (Iba1+; arrows). The lack of Iba1 expression in syncytial cells confirms that they are type II pneumocytes (inset: arrowhead). **E)** T cells (CD3+) are present in moderate numbers throughput the infiltrates and around areas of type II pneumocyte hyperplasia (*) **F)** B cells (CD45R/B220+) form occasional small aggregates in proximity to areas of type II pneumocyte hyperplasia (*). A - artery; B - bronchiole; V - vessel. HE stain; immunohistology, hematoxylin counterstain. Bars represent 20 µm.

In two of the four single SARS-CoV-2 infected and three of the four double infected mice at the later time point (7 days post SARS-CoV-2 infection), we observed a mild or moderate non-suppurative meningoencephalitis, mainly affecting the midbrain and brainstem (Fig. 9). This was more severe in the double infected mice, where the perivascular infiltrates contained degenerate leukocytes and appeared to be associated with focal loss of integrity of the endothelial cell layer (Fig. 9B).

**Figure 9.**
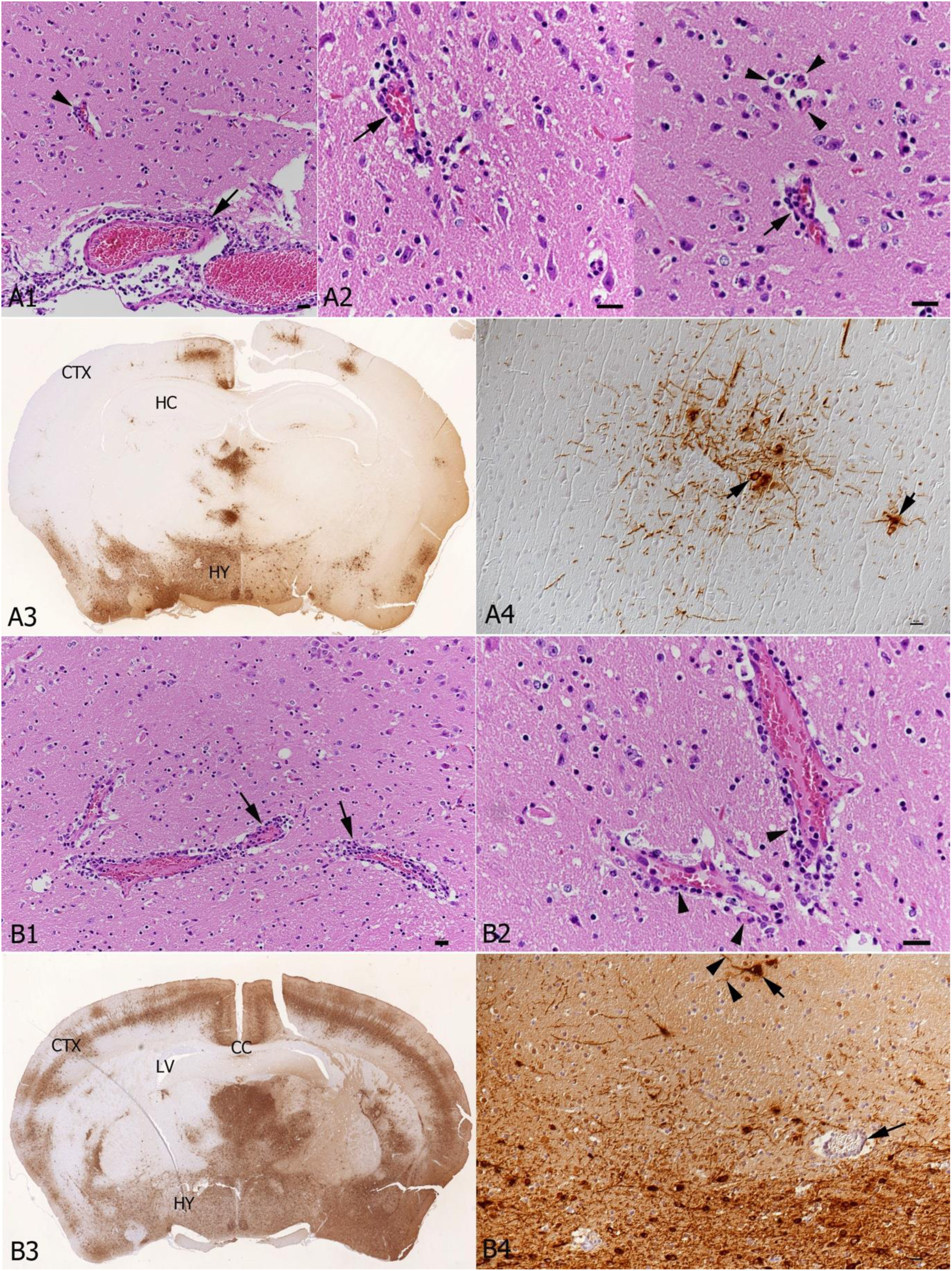
Brain, K18-hACE2 transgenic mice, after day 7 post single infection with SARS-CoV-2 or at day 10 post infection with IAV and day 7 post infection with SARS-CoV-2 in double infections. **A.** SARS-CoV-2 single infection. **A1, A2)** Hypothalamus. **A1)** Vessels in the leptomeninx (arrow) and in the brain parenchyma (arrowhead) exhibit mild perivascular mononuclear infiltrations, consistent with mild non-suppurative meningoencephalitis. **A2)** Higher magnification highlighting the one-layered perivascular infiltrate (arrows). There are a few degenerate cells (right image; arrowheads). **A3)** Coronal section at the level of the hippocampus (HC), showing extensive SARS-CoV-2 antigen expression in the hypothalamus (HY) and bilateral patchy areas with positive neurons also in the cortex (CTX). **A4)** A higher magnification of a focal area with SARS-CoV-2 expression shows that infection is in the neurons (arrowheads). **B)** IAV and SARS-CoV-2 double infected K18-hACE2 transgenic mouse. **B1, B2)** The perivascular mononuclear infiltrate is slightly more intense than in the SARS-CoV-2 single infected mouse (B1: arrows), consistent with a moderate non-suppurative encephalitis. Among the perivascular infiltrate are several degenerate cells (B2: arrowheads). **B3)** Coronal section at the level of the corpus callosum (CC), showing extensive widespread bilateral SARS-CoV-2 antigen expression (HY: hypothalamus; CTX: cortex; LV: left ventricle). HE stain and immunohistology, hematoxylin counterstain. Bars represent 20 µm.

The effect of Fluenz tetra immunisation on the lungs of the mice was assessed by histology and immunohistology. On day 6 after its application, lungs exhibited a mild multifocal increased interstitial cellularity, mild to moderate multifocal mononuclear (macrophages, lymphocytes) peribronchial and perivascular infiltration and mild vascular changes, i.e. mild endothelial cell activation as well as leukocyte rolling and emigration. There were very rare positive intact individual bronchiolar epithelial cells and type II pneumocytes (Supplementary Fig. 3A). SARS-CoV-2 coinfection (3 dpi) was associated with the same histological changes and IAV antigen expression, while SARS-CoV-2 antigen expression was not observed (Supplementary Fig. 3B). Examination of the nasal mucosa at this stage found IAV antigen expression restricted to a few, partly degenerate respiratory epithelial cells in one animal (Supplementary Fig. 3C) while SARS-CoV-2 antigen expression was found in individual and small patches of intact and occasionally degenerate respiratory epithelial cells (Supplementary Fig. 3C). The olfactory epithelium and brain were negative. At day 10 post Fluenz Tetra, histological changes in the lungs were restricted to a mildly increased interstitial cellularity and mild peribronchial and perivascular infiltration by lymphocytes and fewer plasma cells (Supplementary Figure 3D). At this stage, SARS-CoV-2 coinfection (7 dpi) did not result in further changes in two of the four animals; neither showed SARS-CoV-2 NP expression in the lung, but there was evidence of infection, represented by a few individual respiratory epithelial cells in the nasal mucosa that were positive for viral antigen, without viral antigen expression in olfactory epithelium and/or brain. The remaining two animals showed changes consistent with SARS-CoV-2 infection, i.e. focal consolidated areas with several macrophages, activated type II cells and occasional syncytial cells, some lymphocytes and neutrophils, and a few degenerate cells (Supplementary Fig. 3E). SARS-CoV-2 NP expression was seen in a few individual macrophages within the focal lesions and in rare alveoli (type I and II epithelial cells) in one animal, and in several patches of alveoli in the second (Supplemental Fig. 3F). In the first animal, infected epithelial cells were not found in the nasal mucosa, but there were individual neurons and a few larger patches of positive neurons and neuronal processes in the frontal cortex and brain stem. In the second animal, the brain was negative, but there were a few SARS-CoV-2 antigen positive respiratory epithelial cells in the nasal mucosa.

### Distinct transcriptional signatures are associated with infection

The transcriptional profile of lung samples can provide a window on the host response to infection for a respiratory pathogen. Therefore, lung samples were taken at day 6 and day 10 post IAV or Fluenz Tetra infection from all groups of mice (Fig. 1B). Total RNA was purified from cells and both host and viral mRNA (and genomic RNA in the case of SARS-CoV-2) were sequenced on the NovaSeq illumina platform.

Transcripts were counted against the *Mus musculus* transcriptome using Salmon(*33*). Gene counts were inferred through tximport and normalised using the edgeR package before identifying differentially expressed genes using the transcription profile from mock infected mice as the control profile. A total of 22101 genes were identified, where the number of transcripts significantly changing in abundance are presented in Table 1. The top 50 differentially expressed genes form clusters (through the pheatmap hclust parameter) based on the experimental groups, where at day 6 IAV and coinfection belong to the same cluster. By day 10 each experimental group can be distinguished by the transcript expression and are more similar to mock than day 6 mice (Fig 10A). Principle component analysis (PCA) revealed separation between IAV and SARS-CoV-2 groups and overlap between coinfection and IAV infection groups (Fig. 10B). Overlapping signatures are likely representing the non-specific anti-viral response. Contrast matrices were made between mice that were coinfected versus mice that were mock infected and mice that were singly infected (Table 1). When comparing coinfected mice to IAV only at day 6, there were no significant transcripts identified to have changed in abundance, however, by day 10 there were significant differences (Table 1 & Fig. 11A). The coinfected mice had significant changes in comparison to SARS-CoV-2 only infected mice at both day 6 and day 10, where more differences were observed at day 6 (Table 1 & Fig. 11C-D).

**Figure 10:**
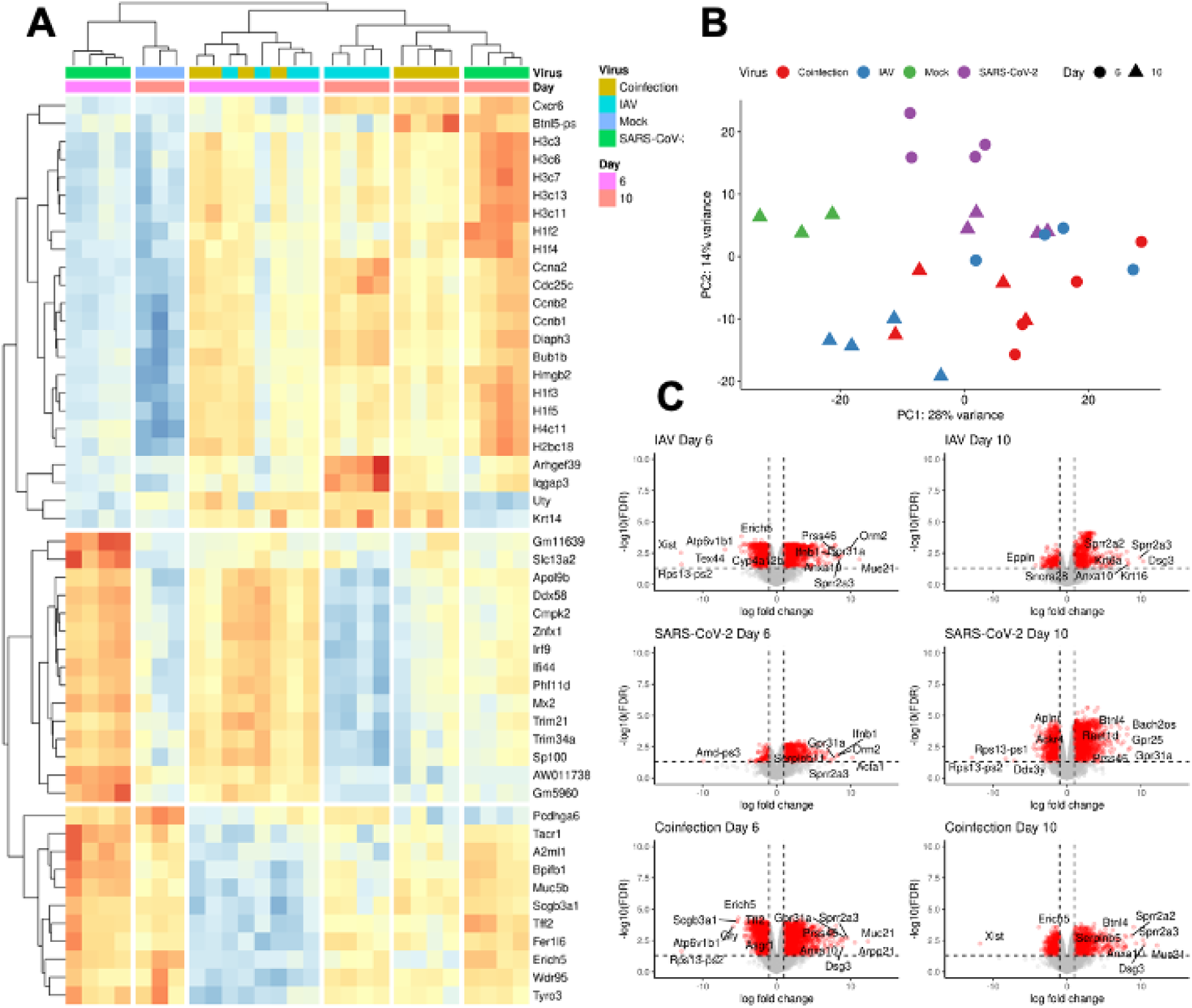
RNA sequencing analysis from hACE2 mice lung homogenates from mice infected with either IAV only, SARS-CoV-2 only or IAV and SARS-CoV-2 (n=3-5). **A)** The top 50 differentially expressed gene transcripts across 4 groups at 2 time-points are shown. **B)** Principal component analysis performed for 28 samples with log2 transformed counts per million (cpm). **C)** Volcano plots comparing differentially expressed genes from each infection group vs mock infected. The horizontal dashed line is representative of a q-value <0.05, and the vertical dashed line is representative of a log2 fold-change of 2. Significant differentially expressed gene transcripts are marked as red.

**Figure 11:**
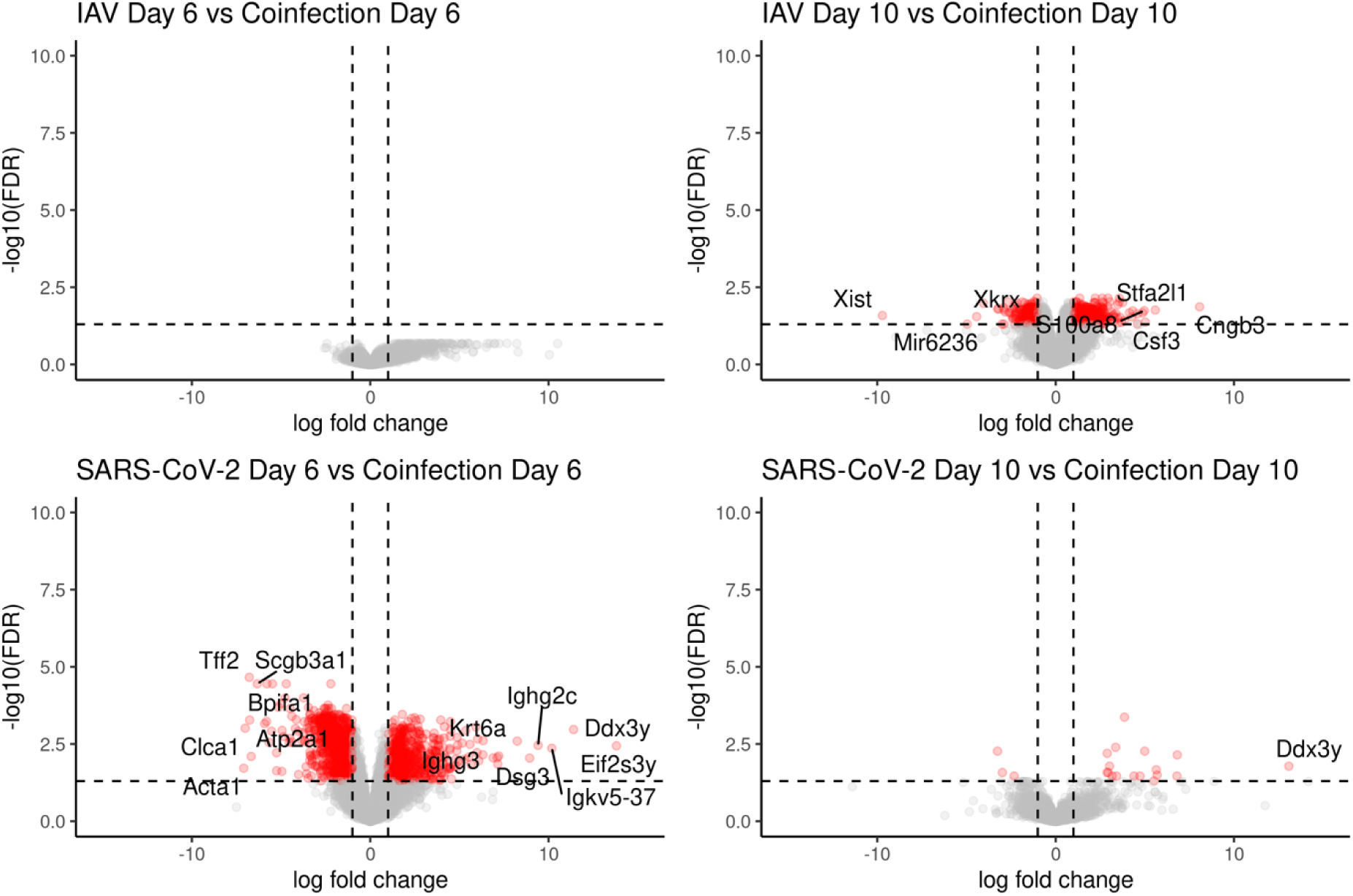
Coinfected mice at day 6 had a similar transcriptomic profile to IAV day 6 infected mice, whereas coinfected mice at day 10 were more similar to SARS-CoV-2 day 10 infected with a few distinct differences. To identify transcripts that were different in each condition, contrasts were made between single infection and coinfection at day 6 and day 10 following a comparison to mock infected. The number of differentially expressed genes are presented in Table 1. There were no significantly changed transcripts when comparing the expression of transcripts in IAV day 6 and coinfection day 6, suggesting these groups were transcriptionally similar.

**Table 1:**
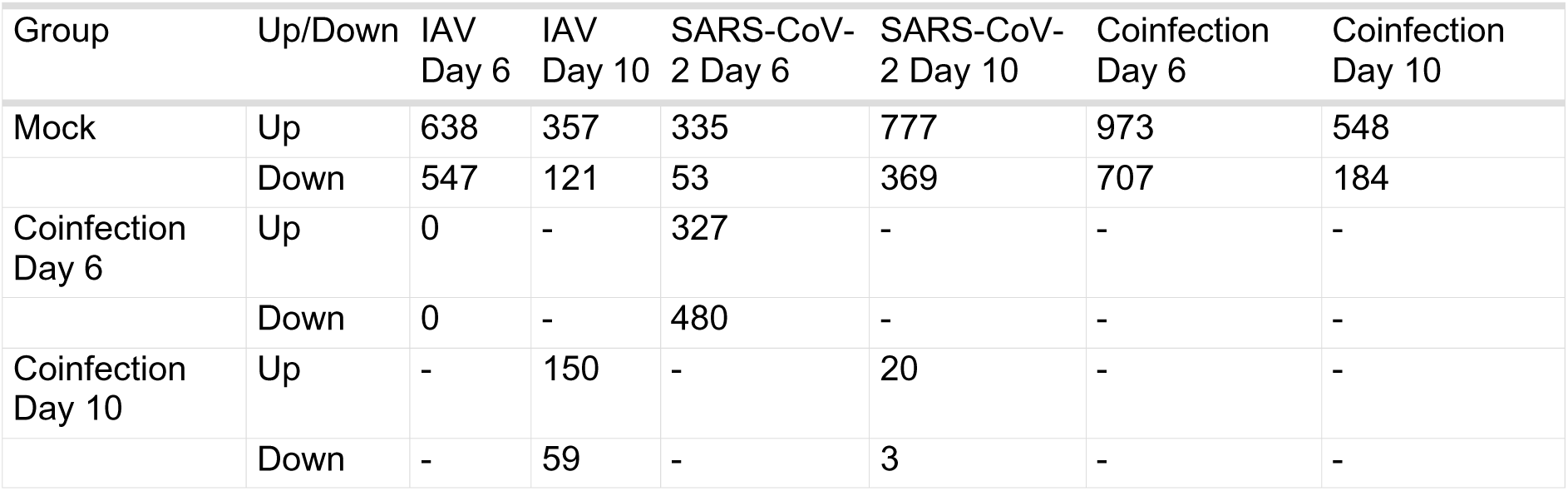
Number of differentially expressed genes with an FDR value less than 0.05 and a log2 fold change more than 2 and less than −2 compared to mock infected mice. Coinfection day 6 and day 10 were compared to day 6 and 10 of individual IAV and SARS-CoV-2 infection.

| Group | Up/Down | IAV Day 6 | IAV Day 10 | SARS-CoV-2 Day 6 | SARS-CoV-2 Day 10 | Coinfection Day 6 | Coinfection Day 10 |
| --- | --- | --- | --- | --- | --- | --- | --- |
| Mock | Up | 638 | 357 | 335 | 777 | 973 | 548 |
|  | Down | 547 | 121 | 53 | 369 | 707 | 184 |
| Coinfection Day 6 | Up | 0 | - | 327 | - | - | - |
|  | Down | 0 | - | 480 | - | - | - |
| Coinfection Day 10 | Up | - | 150 | - | 20 | - | - |
|  | Down | - | 59 | - | 3 | - | - |

To assign ontology to the transcripts identified during differential gene expression analysis, clusterProfiler was used to provide biological process, molecular function and cellular component gene ontology terms. Overall, the terms identified show an upregulation of innate immune responses (Fig.12). By day 10, the IAV group does not show an enrichment for these terms indicative of infection, however, normal cellular processes such as “positive regulation of cell cycle” and “nuclear division” are observed, supporting that this experimental group has resolved the IAV infection. Interestingly, SARS-CoV-2 only infected mice maintain terms representative of an active viral infection, however, do not show enrichment in “leukocyte chemotaxis”, “chemokine production” and “cellular response to interferon gamma” as seen in the coinfected group. Cellular components (Supplementary Fig. 5) and molecular functions (Supplementary Fig. 6) are also presented. Cytokine activity molecular functions are upregulated at day 6 and day 10 in coinfected mice and are otherwise only seen at day 6 in IAV only infected mice.

**Figure 12:**
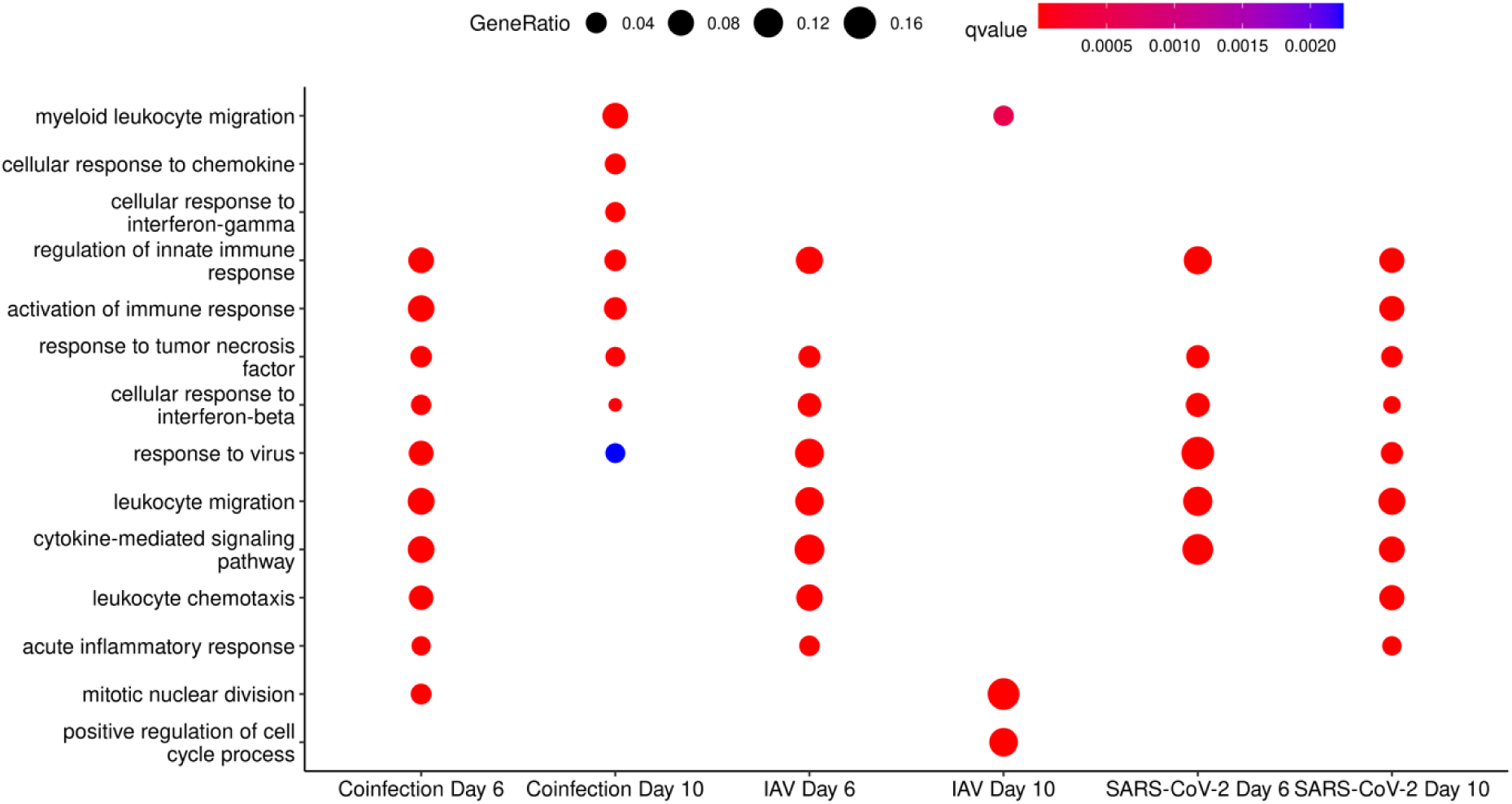
Selected Biological Process GO Terms derived from transcripts in increased abundance when comparing infection groups to mock at Day 6 and Day 10 in clusterProfiler.

Comparing SARS-CoV-2 only and SARS-CoV-2 following a Fluenz Tetra immunisation, a total of 82 significant differentially expressed transcripts were identified at day 6 and a total of 71 at day 10 (Table 2). Interestingly, at day 6 interferon stimulated genes (ISGs) such as Ifit1, Ifit3, and Trim69 were in higher abundance in SARS-CoV-2 infected mice (Supplementary Table 1). As SARS-CoV-2 singly infected mice exhibited respiratory signs and weight loss, it can be postulated that the regulation of these transcripts are important factors for disease severity. By day 10, transcripts associated with adaptive immune responses were found in higher abundance in the Fluenz Tetra immunised SARS-CoV-2 infected mice (Igha, Ighg1, Ighv1-64, Ighv1-75, Ighv5-17, Ighv7-1, Jchain, Igkv13-84, Igkv3-5, Igkv10-96, Igkv12-44, Igkv14-111, Igkv4-57-1) (Supplementary Table 2). As the plethora of transcripts seen to be in higher abundance in Fluenz Tetra immunised mice may offer some protection whilst immunisation seems to dampen the expression of transcripts associated with interferon signalling.

**Table 2:** Number of transcripts changing in abundance and an FDR <= 0.05 when comparing SARS-CoV-2 only infection and coinfection to SARS-CoV-2 infection following Fluenz Tetra immunization at day 6 and day 10.

| Group | Up/Down | SARS-CoV-2<br>Day 6 vs Mock | SARS-CoV-2<br>Day 10 vs Mock | Coinfection<br>Day 6 vs Mock | Coinfection<br>Day 10 vs Mock |
| --- | --- | --- | --- | --- | --- |
| Fluenz Tetra + SARS-CoV-2 Day 6 vs Mock | Up | 57 | - | 416 | - |
|  | Down | 25 | - | 843 | - |
| Fluenz Tetra + SARS-CoV-2 Day 10 vs Mock | Up | - | 17 | - | 108 |
|  | Down | - | 54 | - | 32 |

Transcripts that were found to be increasing and decreasing in such comparison, were inputted into ClusterProfiler to assign gene ontology terms for biological process, molecular function, and cellular components. Biological process clusters demonstrate that the Fluenz Tetra and SARS-CoV-2 infected mice have transcripts associated with “T cell and lymphocyte regulation” in higher abundance than mice infected with SARS-CoV-2 only at day 6 whereas “defence response to virus” and “response to virus” were higher in the SARS-CoV-2 infected only group. By day 10, the SARS-CoV-2 infected group showed an increase in transcripts associated with “leukocyte aggregation”, “leukocyte migration involved in inflammatory response” and “leukocyte chemotaxis”, further demonstrating that SARS-CoV-2 driven pathological processes are driven by host inflammation pathways (Fig. 14).

**Fig 13:**
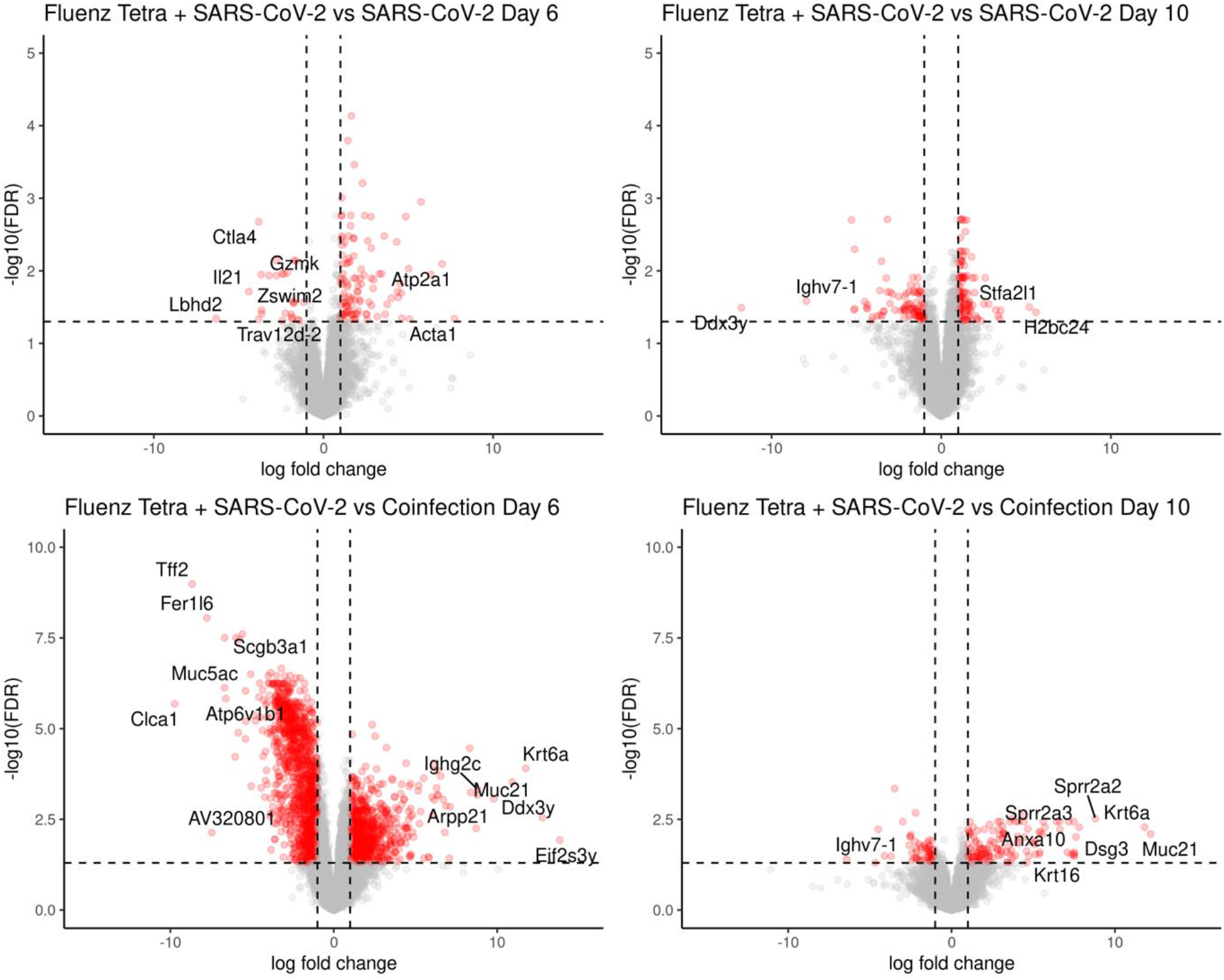
Key differences in transcript abundances between SARS-CoV-2 only infection and coinfection against SARS-CoV-2 infection with the addition of Fluenz Tetra were identified by comparing the two conditions after comparison to mock infected.

**Fig 14:**
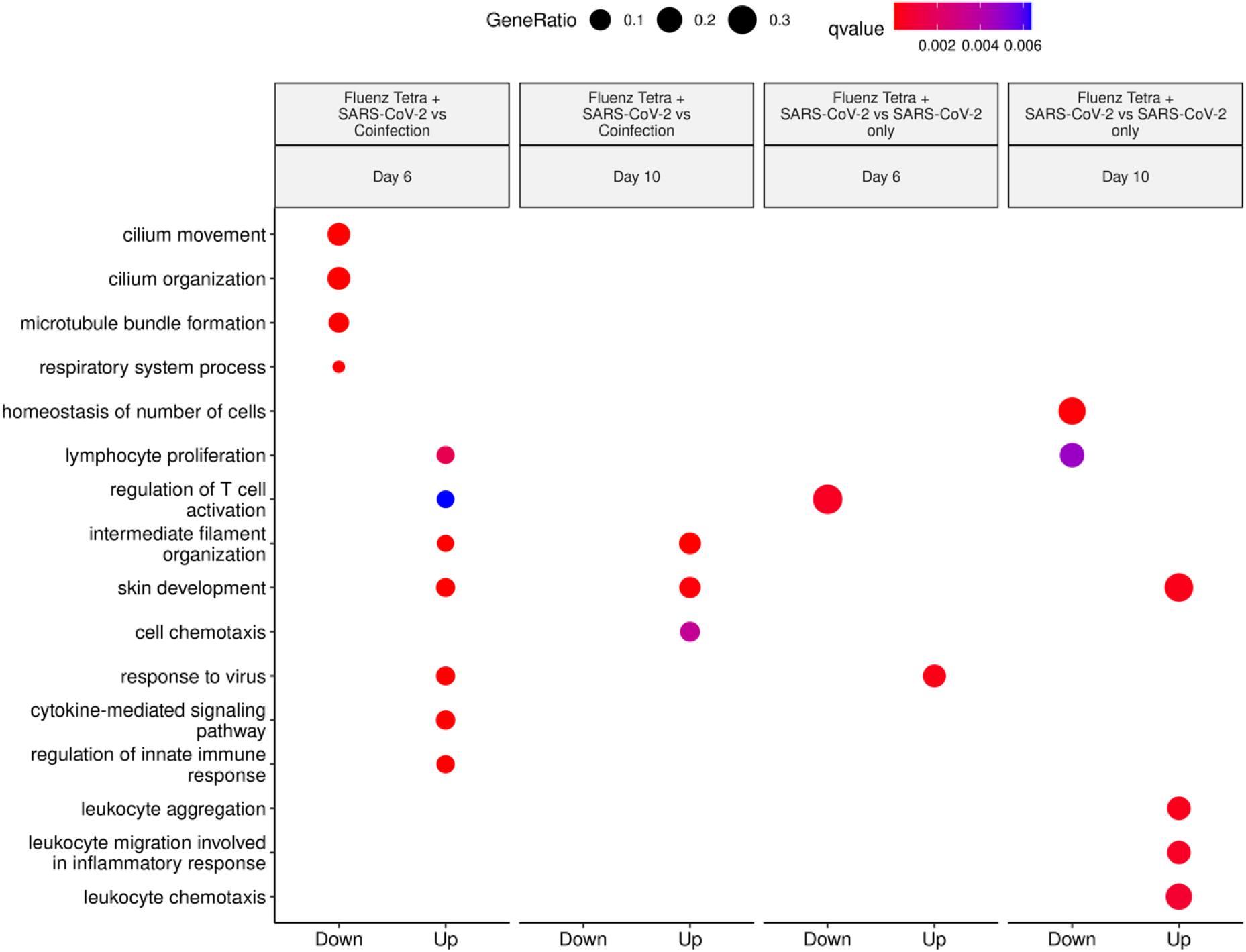
Selected Biological Process GO Terms derived from transcripts increasing and decreasing in abundance when comparing the Fluenz Tetra and SARS-CoV-2 infected group with SARS-CoV-2 only or coinfection groups (following comparison to mock infected). Clusters identified in “up” represent a higher abundance in SARS-CoV-2 infected only or coinfection, whereas “down” represents a higher abundance in Fluenz Tetra and SARS-CoV-2.

When comparing SARS-CoV-2 infection following a Fluenz Tetra immunization to the coinfection group, 1259 differentially expressed genes were identified at day 6, and 140 at day 10 (Table 2). Gene ontology assessment with clusterProfiler revealed enrichment in biological processes such as “cilium movement” and “cilium organisation” in comparison to the coinfected mice. Many genes within this cluster belonged to the cilia and flagella associated protein (cfap) family and are downregulated during infection (Fig. 14). Additionally, processes such as “cytokine production” and “innate immune response” are upregulated in coinfected animals in comparison to the vaccine group, demonstrating that Fluenz Tetra immunisation prevents the transcriptional activity indicative of pulmonary damage.

## Discussion

In this study, sequential infection with IAV followed by SARS-CoV-2 led to more severe pulmonary disease than infections with IAV or SARS-CoV-2 alone. Following IAV infection, mice coinfected with SARS-CoV-2 displayed significantly higher weight loss, elevated respiratory distress and more rapid mortality compared to mice infected with IAV alone. Transcriptomics analysis revealed that the expression of several genes specific to airway epithelial cells such as *Scgb3a1*, C*fap126*, and *Cyp2f2* was more downregulated in coinfected mice compared to singly infected mice at days 6 and 10, which is indicative of increased lung epithelial cell damage. Interestingly, coinfected mice exhibited significantly lower levels of SARS-CoV-2 viral RNA and sgRNA at day 6 (3 dpi with SARS-CoV-2) compared to SARS-CoV-2 singly infected mice, indicating that, whilst coinfection results in enhanced respiratory signs, existing IAV infection interferes with infection of SARS-CoV-2 at this time point. Furthermore, pre-vaccination with an attenuated quadrivalent influenza vaccine (FluenzTetra) also elicited a reduction in SARS-CoV-2 viral RNA and sgRNA, however in the absence of the advanced weight loss, and clinical signs observed in the SARS-CoV-2 IAV coinfected mice. These findings were recapitulated by analysis and comparison of the pathological changes in the lungs. Overt pulmonary damage was due to IAV and was represented by multifocal epithelial cell necrosis in bronchioles and adjacent alveoli. SARS-CoV-2 infection was also multifocal but restricted to alveoli distant from those affected by IAV, consistent with infection via aerosol from the upper respiratory tract and reflecting that both viruses compete for their target cells in alveoli; accordingly, destruction of alveoli by IAV could explain the lower SARS-CoV-2 loads in double infected mice. By day 10, coinfected mice and SARS-CoV-2 singly infected mice displayed similar levels of viral RNA and sgRNA, suggesting that whilst initially inhibited by the presence of IAV, SARS-CoV-2 was able to overcome this inhibition and achieve unconstrained viral replication. In the Fluenz Tetra vaccinated mice SARS-CoV-2, viral RNA and sgRNA copies remained undetectable in 2/4 animals, indicating that the immune response triggered by attenuated influenza virus is restrictive to SARS-CoV-2 infection. This was reflected in the lung transcriptome profile that showed a sustained innate response in coinfected animals over the time period of both infections. Histological examination showed that at 10 days post infection, the damage induced by IAV was resolving. There were distinct regenerative changes, represented by respiratory epithelial cell and type II pneumocyte hyperplasia, accompanied by a moderate macrophage dominated inflammatory response. SARS-CoV-2 infection still showed the same distribution pattern, with viral antigen expression in epithelial cells of unaltered appearing alveoli, but not in hyperplastic type II cells confirming that, like other coronaviruses, also SARS-CoV-2 infects only fully differentiated epithelial cells. However, there was evidence of SARS-CoV-2 induced damage, represented by syncytia formation of type II pneumocytes and more pronounced alveolar damage (acute desquamative pneumonia and occasional hyaline membrane formation). With SARS-CoV-2 single infection the inflammatory response appeared to be less macrophage dominated, as T cells were present in similar amounts. The two Fluenz Tetra vaccinated mice in which SARS-CoV-2 viral RNA and sgRNA was detected also exhibited SARS-CoV-2 associated histological changes and viral antigen expression, though to a lesser degree than the double infected animals, further confirming the limiting effect of influenza immunization on SARS-CoV-2 infection and damage in the lung.

Viral Interference is a well-documented phenomenon which has previously been reported between influenza B viruses (IBV) in a ferret model, in which infection with one IBV subtype was able to prevent infection with another subtype when infections were separated by 3 days (*34*). Similar to the observations described herein between IAV and SARS-CoV-2, coinfection of antigenically unrelated viruses such as IAV and IBV did not confer resistance when challenged within 3 days, but merely delayed the shedding of the challenged virus. This delay in shedding likely accounts for the differing times at which coinfected animals and SARS-CoV-2 singly infected animals lost weight; SARS-CoV-2 singly infected animals exhibited weight loss at 4 dpi, however, coinfected animals began to recover from IAV infection before succumbing to delayed SARS-CoV-2 infection. Unlike this study, wherein IAV and SARS-CoV-2 coinfected animals exhibited significantly increased weight loss, coinfection with IAV and IBV has been reported to lead to delayed viral shedding but did not influence disease severity^28^.

Mathematical modelling and *in vitro* and *in vivo* studies have shown that prior infection with rhinovirus interferes with IAV infection(*6*). This interference is mediated by the induction of interferon stimulated genes (ISGs) following rhinovirus infection which work to suppress IAV. Similarly, infection with IAV results in the activation of the IFN response and the upregulation of ISGs which induce an antiviral state that works to limit infection(*35*) (and reviewed(*36*)). We propose that this response is also active in the K-18hACE2 mice 3 days post IAV infection and contributes to the inhibition of the incoming SARS-CoV-2 infection, thus resulting in lower viral load as measured by RT-qPCR at day 6. IAV viral load was found to be similar between coinfected and IAV singly infected mice, demonstrating that SARS-CoV-2 infection does not interfere with prior IAV infection. Similarly, by day 10 (7 days post SARS-CoV-2 infection) both coinfected and IAV singly infected mice were negative for IAV by immunohistology and qPCR, indicating that SARS-CoV-2 infection does not prolong IAV infection or interfere with the ability of the immune system to clear IAV infection. At this stage, the IAV singly infected mice exhibited histological features consistent with regeneration including hyperplasia of the bronchiolar epithelium and type II pneumocytes. Conversely, while the coinfected mice also displayed evidence of epithelial regeneration, they also presented several hallmarks of acute lung injury including alveolar oedema, fibrin exudation, hyaline membrane formation and degeneration and desquamation of alveolar epithelial cells. This elevated lung injury is consistent with the viral loads of SARS-CoV-2 present in these animals at day 10.

Compared to mice coinfected with IAV and SARS-CoV-2, Fluenz Tetra immunized SARS-CoV-2 infected mice displayed reduced ISG expression at 6 dpi, but elevated expression of transcripts associated with T cell and lymphocyte regulation. The reduction in ISG transcripts in the Fluenz Tetra immunized mice may be due to the low IAV load found in these mice, owing to the attenuation of the virus strains included in the vaccination. Fluenz Tetra does not contain an adjuvant, therefore the observed stimulation of innate and adaptive immune response transcripts is due to viral infection alone (European Medicines Agency, https://www.ema.europa.eu/en/medicines/human/EPAR/fluenz-tetra). Since Fluenz Tetra was able to reduce SARS-CoV-2 viral loads, but without the increase in disease severity associated with IAV infection, it can be postulated that the innate immune response elicited by pathogenic IAV infection contributes to the enhanced disease severity displayed by coinfected animals. As 2/4 Fluenz Tetra vaccinated mice displayed no detectable SARS-CoV-2 viral RNA at days 6 and 10, and no evidence of NP staining in the lungs, this preimmunization appears to have rendered them refractory to SARS-CoV-2 infection.

Interestingly, a similar coinfection study which utilised the IAV strain A/WSN/33 (WSN) found similar pathological findings to this study following coinfection, however they reported that prior infection with IAV enhanced SARS-CoV-2 infection (*37*). IAV infection *in vitro* was found to enhance ACE2 expression, thereby promoting SARS-CoV-2 infection, potentially due to the modulation of ACE2 expression by IFN (*37, 38*). This enhancement was also noted *in vivo* using the K-18 mouse model, however the expression of hACE2 in this model was not upregulated following IAV infection. The observed enhancement of SARS-CoV-2 infection therefore cannot be attributed to the IFN response elicited by WSN infection, and the authors postulate that the observed enhancement of SARS-CoV-2 pathogenicity is a unique feature of IAV infection. The observed differences between these results and the results presented herein may be explained by the doses of virus used and the timings of the infections. The study in which IAV enhanced SARS-CoV-2 infection utilised a higher dose of IAV (2000 PFU) compared to the dose used in our study (100 PFU) and the mice were infected with SARS-CoV-2 a day earlier, at 2 dpi compared to 3 dpi. As IAV is capable of antagonising the IFN response early in infection, this increased dose may result in more potent reduction of the IFN response. Similarly, the early IFN response may be reduced at 2 dpi compared to 3 dpi, thereby promoting SARS-CoV-2 infection. Unfortunately, the authors did not quantify the IFN response in the coinfected mice, nor carry out transcriptomic analysis, so direct comparisons are not possible. Future studies utilising mouse adapted SARS-CoV-2 (*39, 40*) are needed to explore the relationship between SARS-CoV-2 and IAV coinfection in more detail, as the ACE2 expression in these mice will be stimulated more naturally following IAV infection instead of relying on the K-18 promoter.

As has recently been reported in studies using K18-hACE2 transgenic mice(*26, 41, 42*), some of the SARS-CoV-2 singly infected and coinfected mice had developed a non-suppurative meningoencephalitis by day 10, predominantly affecting the midbrain and brainstem. As shown an in-depth study of our group on SARS-CoV-2 associated encephalitis(*42*), the virus spreads to the brain after the initial bout of respiratory infection, usually from day 4 onwards. Distribution of virus antigen and inflammatory changes are consistent with ascending infection from the nasal cavity, via the olfactory bulb(*42, 43*). Interestingly, the coinfected mice displayed more substantial virus spread in the brain and a more pronounced perivascular infiltration with evidence of structural blood-brain-barrier (BBB) breakdown. The mechanism through which coinfection with IAV may enhance SARS-CoV-2 neurological infection is unclear. While brain infection has been well documented in cases of influenza (*44–46*), this is predominantly limited to neurotropic and highly pathogenic strains and occurs via breakdown of the BBB following high levels of viremia (*45, 47*). BBB integrity is also reduced by proinflammatory cytokines such as IL-6, IL-1β and IFN-γ which disrupt the tight junctions maintained by brain microvascular endothelial cells (reviewed in (*48*)). While the IAV X31 strain used herein did not result in brain infection, it is possible that the increased cytokine response present in coinfected animals further compromised the BBB integrity and allowed increased leukocyte recruitment. As brain invasion was only apparent in 1/4 of the Fluenz Tetra immunized mice post SARS-CoV-2 infection, it can be postulated that the innate immune response raised by attenuated IAV was sufficient to reduce SARS-CoV-2 infectivity, without resulting in a pathogenic immune response associated with increased neuroinvasion. The increased pathogenicity associated with coinfection can therefore be the consequence of a pathological overstimulation of the innate immune response.

No animal model can predict with absolute certainty the consequences of coinfection in humans. However, the data presented here may have critical implications for development of successful pre-emptive interventions for SARS-CoV-2. Fortunately, public health interventions aimed at delaying the transmission of SARS-CoV-2 should also provide a consequent reduction in transmission of influenza virus if they are effectively implemented. Moreover, some but not all experimental therapeutics being studied for SARS-CoV-2 have also been demonstrated to exhibit activity against influenza virus. As for other viruses for which successful antiviral interventions have been developed, the SARS-CoV-2 polymerase has emerged as a strong initial target for small molecule inhibitors. Importantly, drugs such as remdesivir and favipiravir that are in various stages of development and clinical evaluation for SARS-CoV-2 have a direct or metabolite-driven *in vitro* activity against influenza virus (*49, 50*), with favipiravir also approved for influenza in Japan. Other examples of dual activity against these viruses are evident with other small molecule antivirals such as nitazoxanide (*51–53*) and niclosamide (*54, 55*), which may present opportunities and/or a basis for prioritisation of candidates for clinical evaluation if necessary exposures can be achieved (*56, 57*). Such antiviral interventions have potential application in treatment of early infection as well as the prophylactic setting. Chemoprevention is a particularly attractive approach when we move into winter months, and selection of the right candidates for evaluation may provide a benefit for both viruses individually and in coinfections. It should be noted that many of the advanced technologies (e.g. broadly neutralising monoclonal antibodies) that are being rapidly accelerated through development have explicit specificities that provide high potency, but this is likely to preclude activity against viruses other than those against which they are directed. The work presented here shows that experimental setting would be an effective pre-clinical platform with which to test therapeutic approaches to dealing with coinfection which is pertinent with the return of seasonal flu outbreaks concomitant with reoccurring global outbreaks of SARS-CoV-2 VOCs.

## Acknowledgements

This work was funded by the US Food and Drug Administration Grant, Characterization of severe coronavirus infection in humans and model systems for medical countermeasure development and evaluation, to JAH, and by the Biotechnology and Biological Sciences Research Council (BBSRC) grants BB/R00904X/1 and BB/R018863/1 to JPS. RPR was supported by a PhD studentship from the MRC Discovery Medicine North (DiMeN) Doctoral Training Partnership (MR/N013840/1). LT is supported by the Wellcome Trust (grant number 205228/Z/16/Z) and the National Institute for Health Research Health Protection Research Unit (NIHR HPRU) in Emerging and Zoonotic Infections (NIHR200907) at University of Liverpool in partnership with Public Health England (PHE), in collaboration with Liverpool School of Tropical Medicine and the University of Oxford. LT is based at University of Liverpool. The views expressed are those of the author(s) and not necessarily those of the NHS, the NIHR, the Department of Health or Public Health England.

## Methods

### Cell culture and virus

Influenza virus A/HKx31 (X31, H3N2) was propagated in the allantoic cavity of 9-day-old embryonated chicken eggs at 35°C for 72 h. Titres were determined by an influenza plaque assay using MDCK cells.

Vero E6 cells (C1008; African green monkey kidney cells) were obtained from Public Health England and maintained in Dulbecco’s minimal essential medium (DMEM) containing 10% foetal bovine serum (FBS) and 0.05 mg/mL gentamycin at 37°C with 5% CO_2_.

UK strain of SARS-CoV-2 (hCoV-2/human/Liverpool/REMRQ0001/2020), which was cultured from a nasopharyngeal swab from a patient, was passaged a further 4 times in Vero E6 cells (*30*). The fourth passage of virus was cultured MOI of 0.001 in Vero E6 cells with DMEM containing 4% FBS and 0.05 mg/mL gentamycin at 37°C with 5% CO_2_ and was harvested 48 h post inoculation. Virus stocks were stored at −80°C. The intracellular viral genome sequence and the titre of virus in the supernatant were determined. Direct RNA sequencing was performed as describe previously (*31*) and an inhouse script was used to check for deletions in the mapped reads. The Illumina reads were mapped to the England/2/2020 genome using HISAT and the consensus genome was called using an in-house script based on the dominant nucleotide at each location on the genome. The sequence has been submitted to Genbank, accession number MW041156. Fluenz tetra (AstraZeneca) is a commercially available quadrivalent live vaccine composed of attenuated A/Victoria/2570/2019 (H1N1) pdm09, A/Cambodia/e0826360/2020 (H3N2), B/Washington/02/2019 and B/Phuket/3073/2013.

### Ethics and clinical information

The patient from which the virus sample was obtained gave informed consent and was recruited under the International Severe Acute Respiratory and emerging Infection Consortium (ISARIC) Clinical Characterisation Protocol CCP (https://isaric.net/ccp), reviewed and approved by the national research ethics service, Oxford (13/SC/0149). Samples from clinical specimens were processed at containment level 3 at the University of Liverpool.

#### Biosafety

All work was performed in accordance with risk assessments and standard operating procedures approved by the University of Liverpool Biohazards Sub-Committee and by the UK Health and Safety Executive. Work with SARS-CoV-2 was performed at containment level 3 by personnel equipped with respirator airstream units with filtered air supply.

### Mice

Animal work was approved by the local University of Liverpool Animal Welfare and Ethical Review Body and performed under UK Home Office Project Licence PP4715265. Mice carrying the human ACE2 gene under the control of the keratin 18 promoter (K18-hACE2; formally B6.Cg-Tg(K18-ACE2)2Prlmn/J) were purchased from Jackson Laboratories. Mice were maintained under SPF barrier conditions in individually ventilated cages.

### Virus infection

Animals were randomly assigned into multiple cohorts. For IAV infection, mice were anaesthetized lightly with KETASET i.m. and inoculated intra-nasally with 10^2^ PFU IAV X31 in 50 µl sterile PBS. For SARS-CoV-2 infection, mice were anaesthetized lightly with isoflurane and inoculated intra-nasally with 50 µl containing 10^4^ PFU SARS-CoV-2 in PBS. For Fluenz Tetra immunization, mice were anesthatised with isoflurane and intranasally inoculated with 50 μl of vaccine formulation. Each 50ul of Fluenza Tetra contains around 2×10^6±4^ of A/Victoria/2570/2019 (H1N1)pdm09, A/Cambodia/e0826360/2020 (H3N2), B/Washington/02/2019 and B/Phuket/3073/2013. Mice were sacrificed at variable time points after infection by an overdose of pentabarbitone. Tissues were removed immediately for downstream processing.

### Histology, immunohistology

For SARS-CoV-2 and IAV coinfection studies, the left lung, kidneys, liver and brain as well as the head were fixed in 10% neutral buffered formalin for 24-48 h and routinely paraffin wax embedded (prior to embedding, the heads were sawn in the midline and gently decalcified in RDF (Biosystems) for twice 5 days, at room temperature (RT) and on a shaker). Consecutive sections (3-5 µm) were either stained with hematoxylin and eosin (HE) or used for immunohistology (IH). IH was performed to detect viral antigens and ACE2 expression and to identify macrophages, T cells and B cells using the horseradish peroxidase (HRP) method. The following primary antibodies were applied: rabbit anti-human ACE2 (Novus Biologicals; clone SN0754; NBP2-67692), goat anti-IAV (Meridian Life Sciences Inc., B65141G), rabbit anti-Iba-1 (antigen: AIF1; Wako Chemicals; macrophage marker), mAB rabbit anti-mouse CD3 (clone SP7: Spring Bioscience, Ventana Medical Systems, Tucson, USA; T cell marker) and rat anti-mouse CD45R (clone B220, BD Biosciences; B cell marker), following previously published protocols(*58–60*), and rabbit anti-SARS-CoV nucleocapsid protein (Rockland, 200-402-A50). Briefly, after deparaffination, sections underwent antigen retrieval in citrate buffer (pH 6.0; Agilent) for ACE2 and SARS-CoV-2 NP detection, and Tris/EDTA buffer (pH 9) for IAV for 20 min at 98 °C, followed by incubation with the primary antibodies (diluted in dilution buffer, Agilent; anti-IAV 1:200, anti-ACE2, 1:200 and anti-SARS-CoV 1:3000) and overnight at 4⁰C for ACE2 and SARS-CoV-2 and for 1 h at RT for IAV. This was followed by blocking of endogenous peroxidase (peroxidase block, Agilent) for 10 min at RT and incubation with the secondary antibodies, EnVision+/HRP, Rabbit (Agilent) for ACE2 and SARS-CoV, and rabbit anti-goat Ig/HRP (Agilent) for IAV), for 30 min at RT, and EnVision FLEX DAB+ Chromogen in Substrate buffer (Agilent) for 10 min at RT, all in an autostainer (Dako). Sections were subsequently counterstained with hematoxylin.

In addition, four mock infected transgenic control mice were subjected to a phenotyping, examining upper and lower airways, lungs, kidneys, liver, small intestine, spleen and brain for any histological changes and for ACE2 expression. Lungs and brain from an age-matched wild type BALB-C mouse stained for ACE2 served to assess any differences in the ACE expression pattern in the transgenic mice, and two wild type C57BL6/J mice infected intranasally with 10^2^ PFU IAV X31 in 50 µl sterile PBS and sacrificed at 6 days post infection served to assess any effect of hACE2 expression in the course of IAV infection. A formalin-fixed, paraffin embedded cell pellet infected with IAV for 24 h served as positive control for the IAV staining, and the spleen of a control mouse as positive control for the leukocyte markers.

For the Fluenz Tetra immunized mice, the left lung and head were fixed in 10% neutral buffered formalin for 24-48 h. Heads were then sawn (coronal sections) with a diamond saw (Exakt 300; Exakt) to prepare appr. 1.5 mm thick tissue slices from apical nose to the level of the atlanto-occipital joint. Heads were gently decalcified in RDF (Biosystems) for twice 5 days, at RT and on a shaker. Heads, including brains and lungs were routinely paraffin wax embedded and stained as described above.

### RNA extraction and DNase treatment

The upper lobes of the right lung were dissected and homogenised in 1ml of TRIzol reagent (Thermofisher) using a Bead Ruptor 24 (Omni International) at 2 meters per second for 30 sec. The homogenates were clarified by centrifugation at 12,000xg for 5 min before full RNA extraction was carried out according to manufacturer’s instructions. RNA was quantified and quality assessed using a Nanodrop (Thermofisher) before a total of 1 μg was DNase treated using the TURBO DNA-free™ Kit (Thermofisher) as per manufacturer’s instructions.

### qRT-PCR for viral load

Viral loads were quantified using the GoTaq® Probe 1-Step RT-qPCR System (Promega). For quantification of SARS-COV-2 the nCOV_N1 primer/probe mix from the SARS-CoV-2 (2019-nCoV) CDC qPCR Probe Assay (IDT) were utilised while the standard curve was generated via 10-fold serial dilution of the 2019-nCoV_N_Positive Control (IDT) from 10^6^ to 0.1 copies/reaction. The E sgRNA primers and probe have been previously described (https://www.nature.com/articles/s41586-020-2196-x) and were utilised at 400nM and 200nM respectively. Murine 18S primers and probe sequences ere utilsied at 400nM and 200nM respectively. The IAV primers and probe sequences are published as part of the CDC IAV detection kit (20403211). The IAV reverse genetics plasmid encoding the NS segment was diluted 10-fold from 10^6^ to 0.1 copies/reaction to serve as a standard curve. The thermal cycling conditions for all qRT-PCR reactions were as follows: 1 cycle of 45°C for 15 min and 1 cycle of 95°C followed by 40 cycles of 95°C for 15 sec and 60°C for 1 minute. The 18s standard was generated by the amplification of a fragment of the murine 18S cDNA using the primers F: ACCTGGTTGATCCTGCCAGGTAGC and R: GCATGCCAGAGTCTCGTTCG. Similarly, the E sgRNA standard was generated by PCR using the qPCR primers. cDNA was generated using the SuperScript IV reverse transcriptase kit (Thermofisher) and PCR carried out using Q5® High-Fidelity 2X Master Mix (New England Biolabs) as per manufacturer’s instructions. Both PCR products were purified using the QIAquick PCR Purification Kit (Qiagen) and serially diluted 10-fold from 10^10^ to 10^4^ copies/reaction to form the standard curve.

### Illumina RNA seq

Following the manufactures protocols, total RNA from lung tissue were used as input material in to the QIAseq FastSelect –rRNA HMR (Qiagen) protocol to remove cytoplasmic and mitochondrial rRNA with a fragmentation time of 7 or 15 minutes. Subsequently the NEBNext® Ultra™ II Directional RNA Library Prep Kit for Illumina® (New England Biolabs) was used to generate the RNA libraries, followed by 11 cycles of amplification and purification using AMPure XP beads. Each library was quantified using Qubit and the size distribution assessed using the Agilent 2100 Bioanalyser and the final libraries were pooled in equimolar ratios. The raw fastq files (2 × 150 bp) generated by an Illumina® NovaSeq 6000 (Illumina®, San Diego, USA) were trimmed to remove Illumina adapter sequences using Cutadapt v1.2. (Cut adapt reference). The option “−O 3” was set, so the that 3’ end of any reads which matched the adapter sequence with greater than 3 bp was trimmed off. The reads were further trimmed to remove low quality bases, using Sickle v1.200 (https://github.com/najoshi/sickle) with a (*61*) minimum window quality score of 20. After trimming, reads shorter than 10 bp were removed.

### RNA sequencing bioinformatic analysis

Trimmed paired end sequencing reads were inputted into salmon (v1.5.2) using the -l A –validateMappings –SeqBias –gcBias parameters. Quant files generated with salmon were imported into RStudio (4.1.1) using tximport to infer gene expression (*62*). The edgeR package (3.34.1) was used to normalise sequencing libraries and identify differentially expressed genes, defined as at least a 2-fold difference from the mock infected group and a false discovery rate (FDR) less than 0.05 51. Principle component Analysis (PCA), volcano plots, heatmaps and Venn diagrams were produced in R studio using the following packages: edgeR, ggplot2 (3.3.5) and pheatmap (1.0.12). Differential gene expression data was used for gene ontology enrichment analysis of biological process terms in each group using the compareCluster function with enrichGO in the ClusterProfiler package (4.0.5) programme in R (*63*). Code used to analyse data is available at https://github.com/Hiscox-lab/k18-hACE2-coinfection-transcriptomics. Sequencing reads are available under BioProject ID: PRJNA886870 on Short Read Archive (SRA).

### Statistical analysis

Data were analysed using the Prism package (version 5.04 Graphpad Software). *P* values were set at 95% confidence interval. A repeated-measures two-way ANOVA (Bonferroni post-test) was used for time-courses of weight loss; two-way ANOVA (Bonferroni post-test) was used for other time-courses; log-rank (Mantel-Cox) test was used for survival curves. All differences not specifically stated to be significant were not significant (p > 0.05). For all figures, *p < 0.05, **p <0.01, ***p < 0.001.

## Supplementary figures

**Figure S1.**
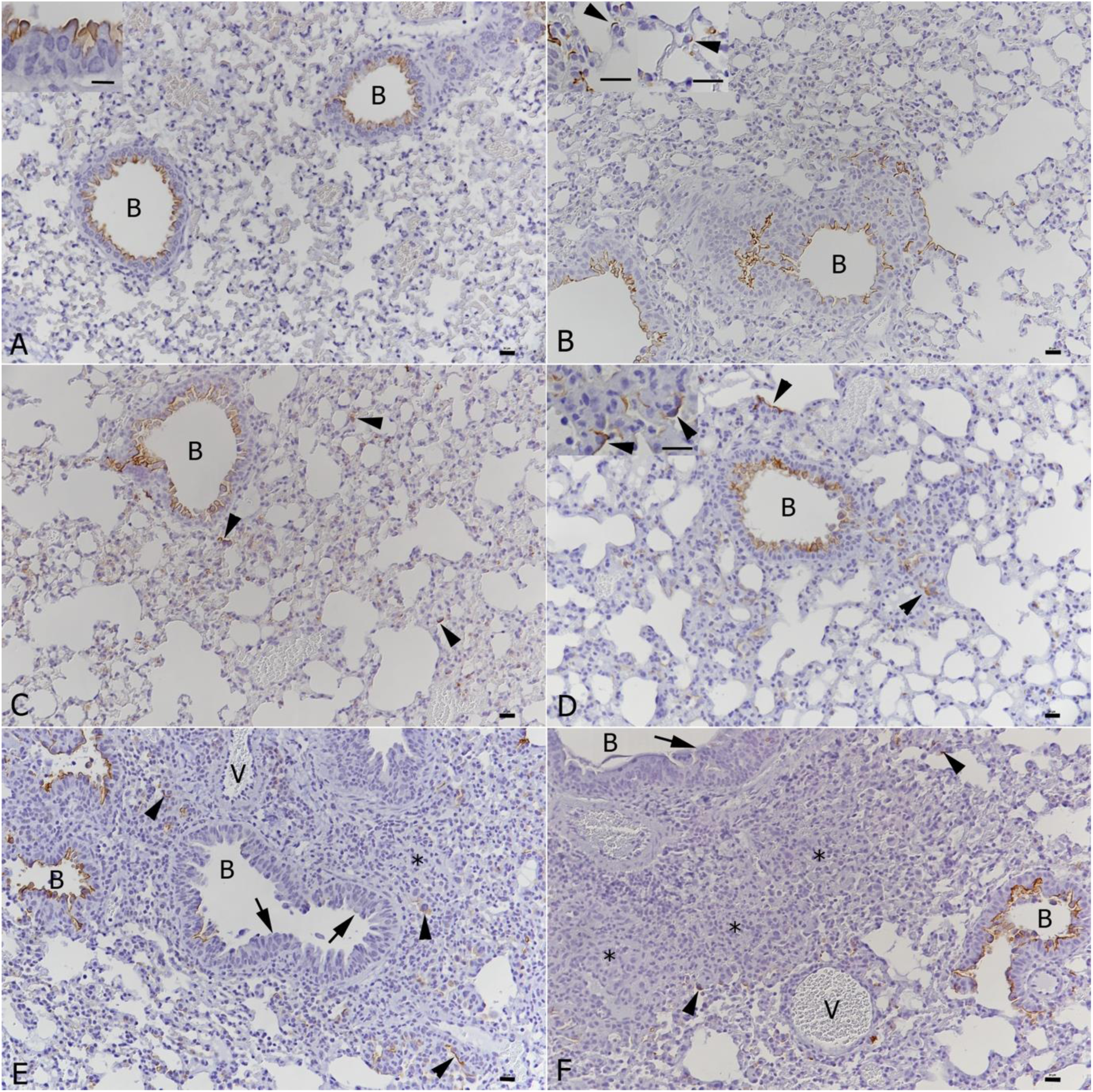
Lung. ACE2 expression pattern in mock infected wild type and K18-hACE2 transgenic mice, and in K18-hACE2 transgenic mice after SARS-CoV-2 and IAV single and double infection. **A**. Wildtype mouse, mock infected. ACE2 expression is seen in bronchiolar (B) epithelial cells; inset: expression is seen along the luminal surface. **B**. K18-hACE2 transgenic mouse, mock infected. ACE2 expression is seen in bronchiolar (B) epithelial cells and in scattered type II pneumocytes (insets). **C**. K18-hACE2 transgenic mouse, 3 days after SARS-CoV-2 infection. The ACE2 expression pattern is identical to that observed in mock infected mice (B: bronchiole; arrowheads: ACE2-positive type II pneumocytes). **D**. K18-hACE2 transgenic mouse, 7 days after SARS-CoV-2 infection. The ACE2 expression pattern is identical to that observed in mock infected mice and mice at 3 days post infection. The amount of ACE2-positive type II pneumocytes is increased; the cells appear activated (inset: arrowheads). **E**. K18-hACE2 transgenic mouse, 10 days after IAV infection. There is a mild increase in ACE2 positive type II pneumocytes (arrowheads). Interestingly, both hyperplastic epithelial cells in the bronchioles (B; arrows) and hyperplastic type II pneumocytes (*) do not express ACE2. **F**. K18-hACE2 transgenic mouse, IAV (10 dpi) and SARS-CoV-2 (7 dpi) double infection. The hACE2 expression pattern is similar to that seen after single IAV infection at 10 dpi. ACE2 expression lacks in hyperplastic bronchiolar (B) epithelial cells (arrow) and hyperplastic type II pneumocytes (*). There are a few individual positive type II pneumocytes (arrowheads). V – vessel. HE stain and immunhistology, hematoxylin counterstain. Bars represent 20 µm.

**Figure S2.**
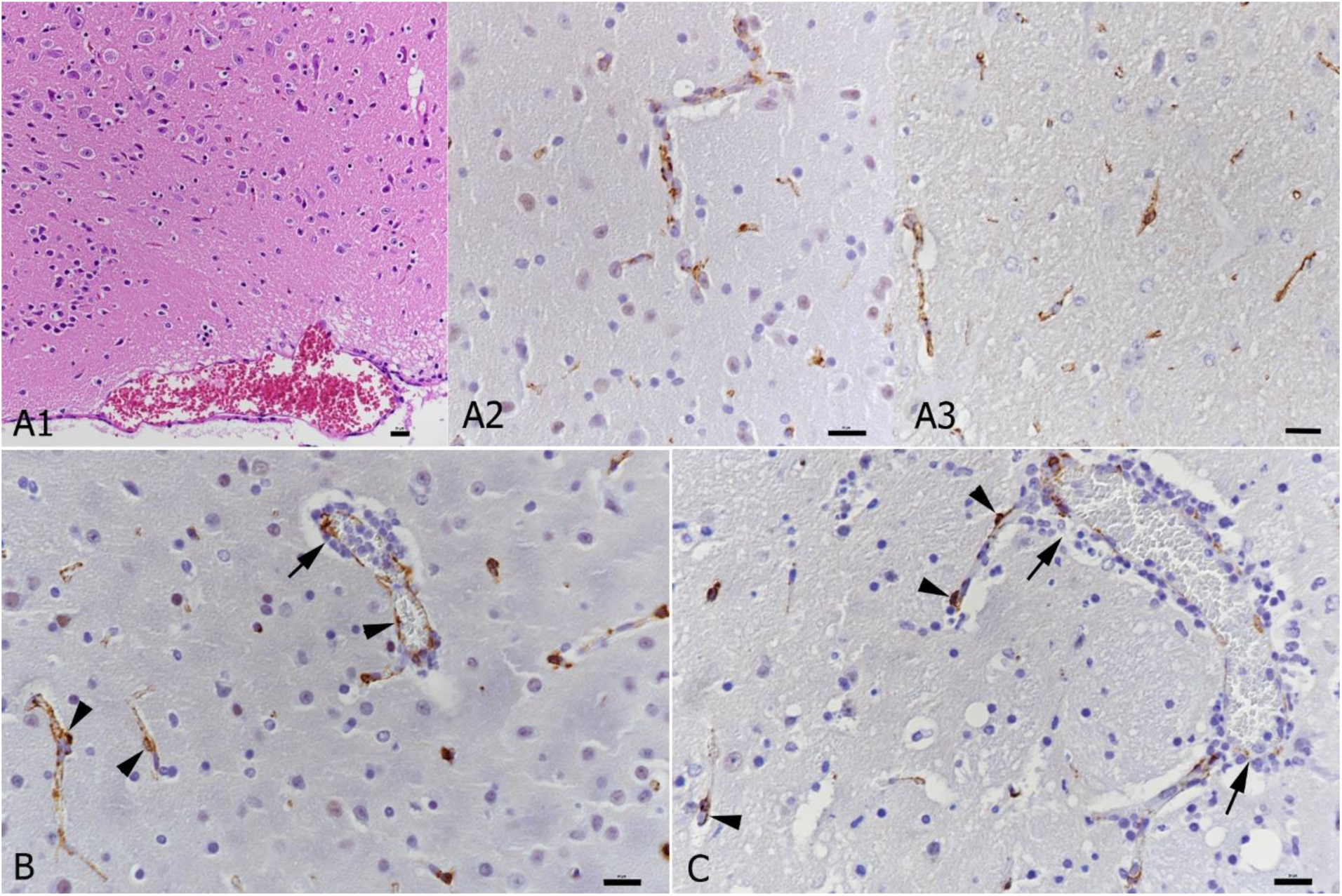
ACE2 expression in the brain. **A.** Mock infected control animals. **A1**. K18-hACE2 transgenic mouse, unaltered hypothalamus. **A2, A3.** ACE2 expression in the microvasculature in a K18-hACE2 transgenic mouse (A2) and wildtype mouse (A3). **B.** SARS-CoV-2 infected K18-hACE2 transgenic mouse. Staining for ACE2 indicates an intact capillary wall (arrowheads) also in areas of perivascular infiltration (arrow). **C.** IAV and SARS-CoV-2 double infected K18-hACE2 transgenic mouse. Staining for ACE2 (arrowheads) is lost in areas of more intense infiltration (arrows). HE stain and immunohistology, hematoxylin counterstain. Bars represent 20 µm.

**Figure S3.**
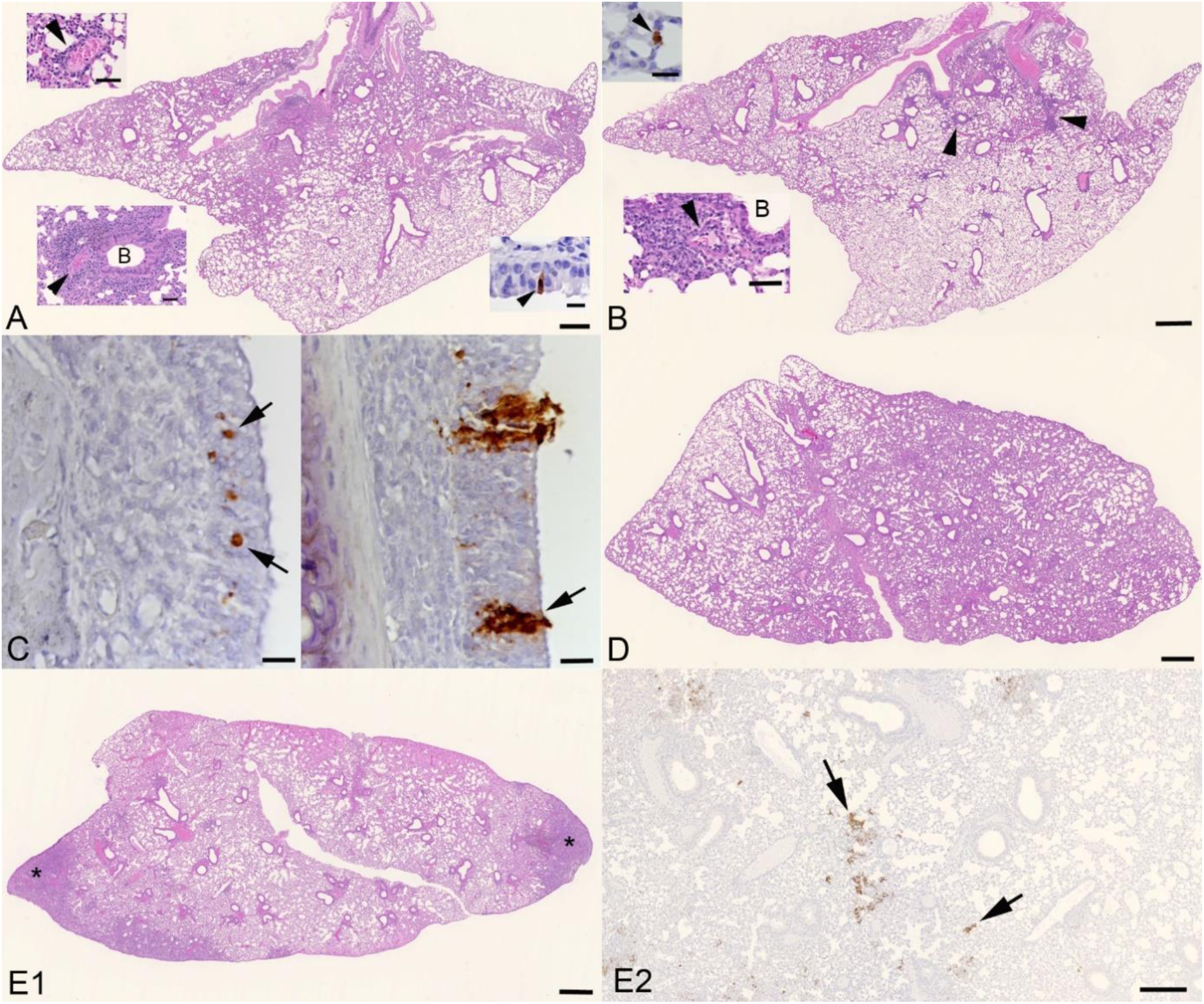
Lung and nose in K18-hACE2 mice after immunization with Fluenz tetra, after immunization and subsequent intranasal infection with SARS-CoV-2, or with SARS-CoV-2 infection alone. **A**. Lung at 6 days post Fluenz Tetra immunization. There is a mild multifocal increase in interstitial cellularity and mild multifocal mononuclear peribronchial (B: bronchiole) and perivascular (arrowhead) infiltration (bottom left inset) and vasculitis (arrowhead; top left inset). IAV antigen expression is seen in rare bronchiolar epithelial cells (arrowhead; right inset) and pneumocytes. HE stain and immunohistology; bars represent 500 µm and 50 µm (insets). **B**. Lung at 6 days post Fluenz Tetra immunization and 3 dpi with SARS-CoV-2. There is a mild multifocal increase in interstitial cellularity and mild multifocal mononuclear peribronchial and perivascular infiltration (arrowheads) and vasculitis (arrowhead; bottom inset). Staining for IAV antigen shows an individual positive type II pneumocyte (arrowhead; top inset). HE stain and immunohistology; bars represent 500 µm, 20 µm (top inset) and 50 µm (bottom inset). **C**. Nose at 6 days post Fluenz Tetra immunization and 3 dpi with SARS-CoV-2. Left: Respiratory epithelium with a few individual IAV antigen-positive cells (arrows). Right: Respiratory epithelium with patches of SARS-CoV-2 antigen-positive cells (arrow). Immunohistology; bars = 20 µm. **D**. Lung at 10 days post Fluenz Tetra immunization. Apart from a very mild increase in interstitial cellularity and very mild mononuclear peribronchial and perivascular infiltration, the lung parenchyma is unaltered. HE stain; bar represents 500 µm. **E**. Lung at 10 days post Fluenz Tetra immunization and 7 dpi with SARS-CoV-2. **E1**. In addition to a mild increase in interstitial cellularity and mononuclear peribronchial and perivascular infiltration there are several focal subpleural areas of consolidation (*) with activated and occasional syncytial type II pneumocytes and infiltration by macrophages, some lymphocytes and neutrophils, with some degenerate cells. HE stain; bar represents 500 µm. **E2**. There are several patches of alveoli with SARS-CoV-2 antigen positive type I and II pneumocytes (arrows). Immunohistology; bar represents 250 µm.

**S5:**
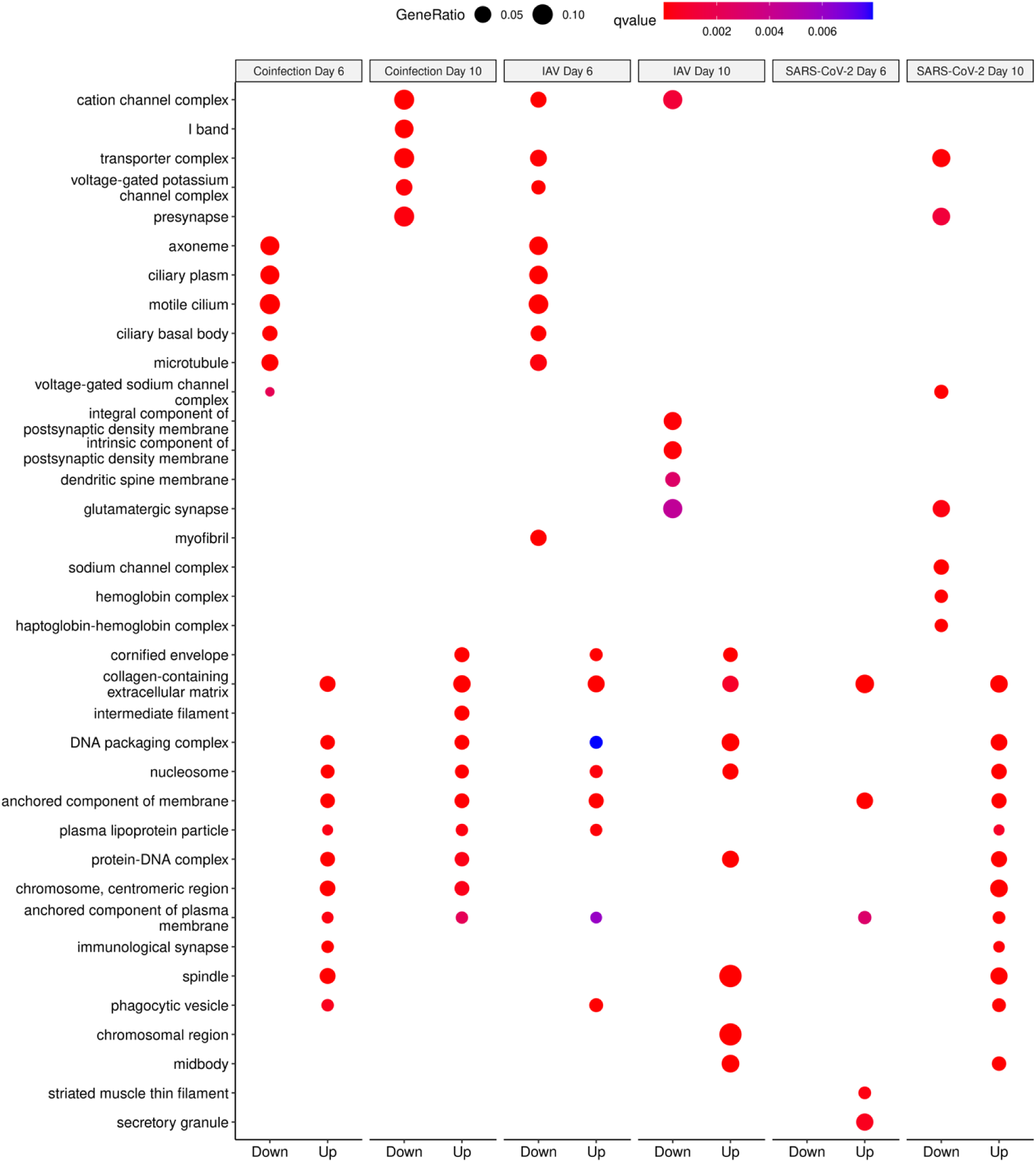
The top 10 cellular component terms reported from clusterProfiler to assess gene enrichment following differential gene expression analysis.

**S6:**
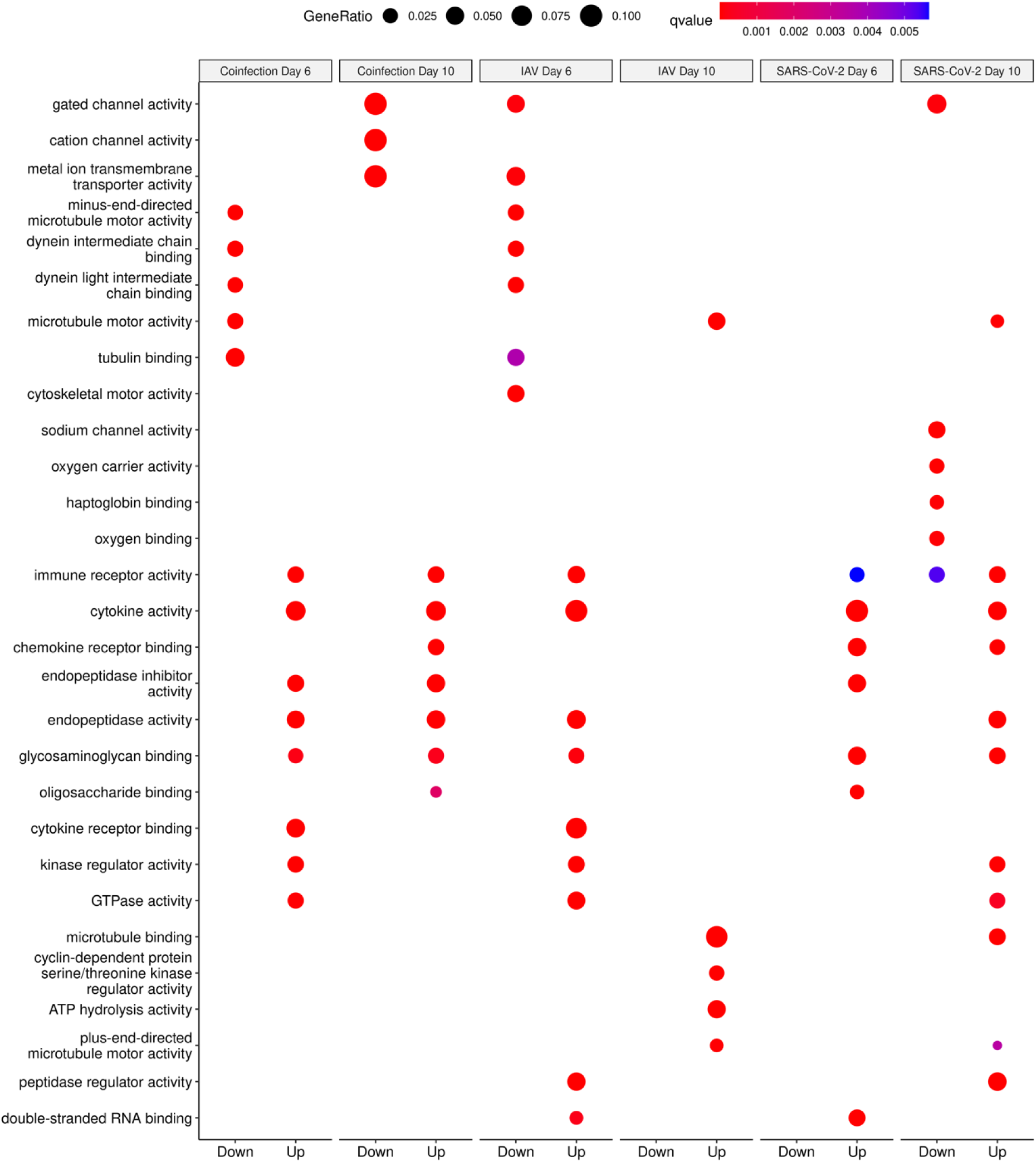
The top 10 molecular function terms reported from clusterProfiler to assess gene enrichment following differential gene expression analysis.

**Figure S7:**
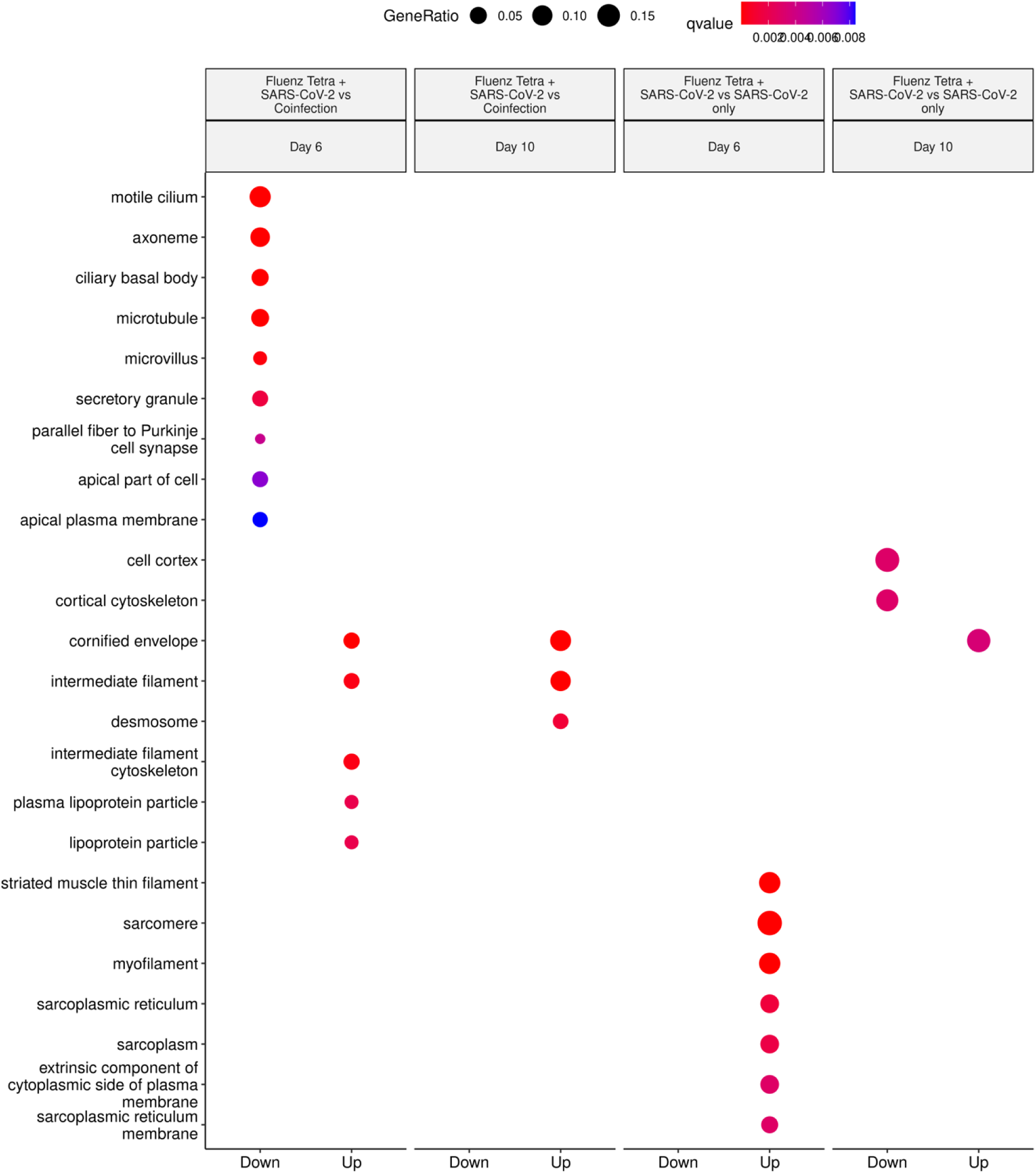
Cellular component GO Terms derived from transcripts increasing and decreasing in abundance when comparing the Fluenz tetra and SARS-CoV-2 infected group and SARS-CoV-2 only infected group (following comparison to mock infected). Clusters identified in “up” represent a higher abundance in SARS-CoV-2 infected only, whereas “down” represents a higher abundance in Fluenz tetra and SARS-CoV-2.

**Figure S8:**
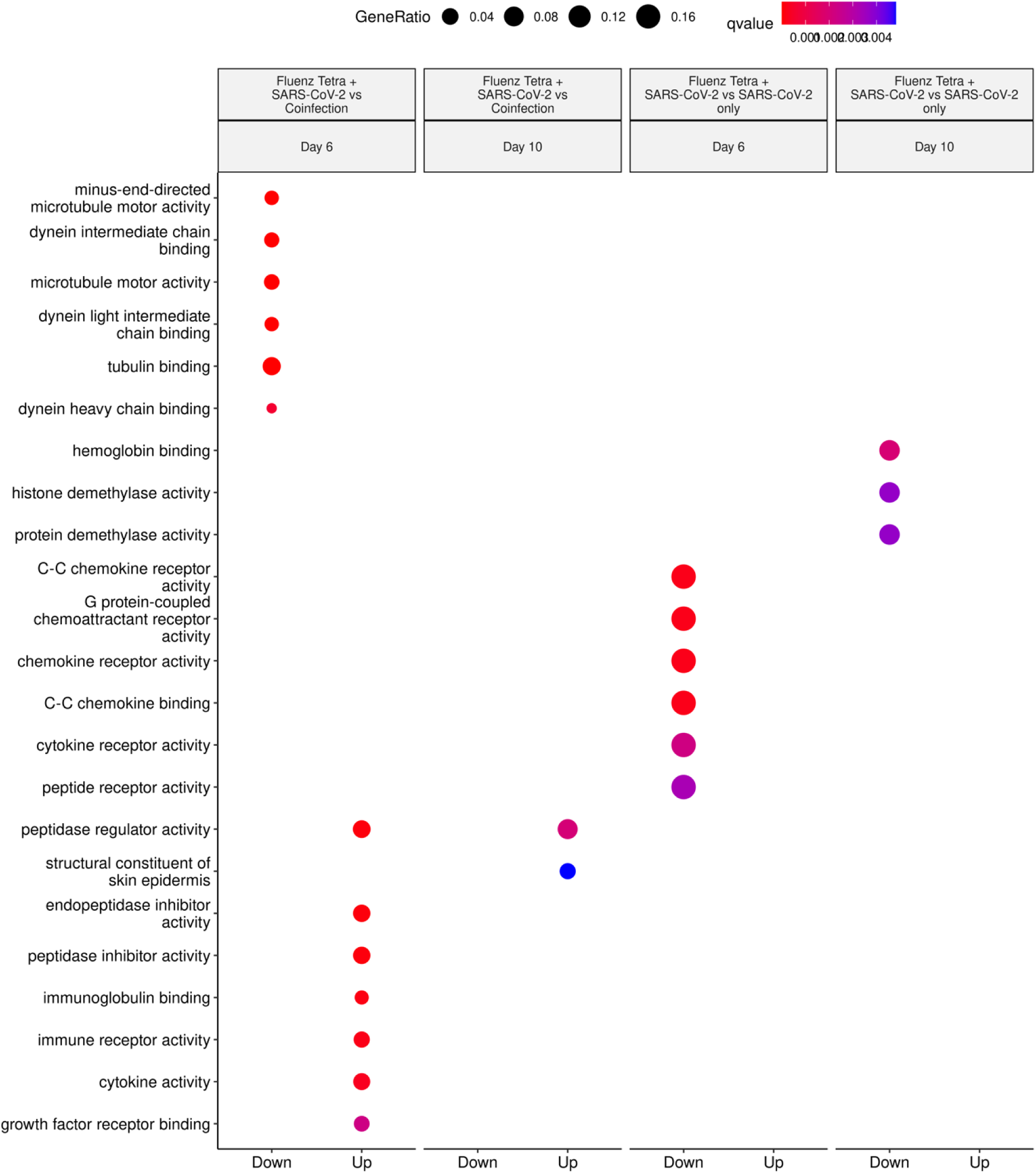
Molecular Function GO Terms derived from transcripts increasing and decreasing in abundance when comparing the Fluenz tetra and SARS-CoV-2 infected group and SARS-CoV-2 only infected group (following comparison to mock infected). Clusters identified in “up” represent a higher abundance in SARS-CoV-2 infected only, whereas “down” represents a higher abundance in Fluenz tetra and SARS-CoV-2.

